# Visualizing the M2 muscarinic acetylcholine receptor activation regulated by aromatic ring dynamics

**DOI:** 10.1101/2024.02.18.580854

**Authors:** Zhou Gong, Xu Zhang, Maili Liu, Changwen Jin, Yunfei Hu

**Affiliations:** State Key Laboratory of Magnetic Resonance and Atomic and Molecular Physics, National Center for Magnetic Resonance in Wuhan, Innovation Academy for Precision Measurement Science and Technology, Chinese Academy of Sciences, Wuhan, 430071, China; Joint Laboratory of the National Centers for Magnetic Resonance in Wuhan and in Beijing, Wuhan, 430071, China; Beijing Nuclear Magnetic Resonance Center, College of Chemistry and Molecular Engineering and Beijing National Laboratory for Molecular Sciences, Peking University, Beijing, 100871, China

## Abstract

A detailed molecular understanding of the G-protein coupled receptor (GPCR) activation mechanism is crucial for rational drug design. Despite the growing number of GPCR structures being resolved in the inactive and activated states, the detailed molecular mechanism of how receptors transits from the inactive towards the active one upon agonist binding remains to be further understood. Herein, we performed comprehensive atomic-level simulations of the M2 muscarinic receptor (M2R) to determine how ligand binding modulates the receptor conformational dynamics. The results reveal that aromatic residue dynamics are closely associated with receptor activation. Binding of antagonists or agonists differentially alters the interacting patterns of critical aromatic residues, stabilizing them into distinct conformations. In addition, we found that the change of interaction and dynamics of the W400^6.48^-F396^6.44^ pair at the transmembrane core plays an essential role in the structural transition of M2R from the inactive state into an active-like state induced by binding of the supra-physiological agonist iperoxo. Moreover, we found that the sidechain dynamics of Y206^5.58^ is important in modulating the conformation of the intracellular cavity. Our work highlights the role of protein conformational dynamics in M2R activation, and provides new insights for future drug designs.

## INTRODUCTION

G-protein coupled receptors (GPCRs) are the largest family of transmembrane proteins that mediate cell signaling in many physiological processes ^1^. Ligand binding to the extracellular pocket induces receptor conformational rearrangements that propagate through the transmembrane (TM) core into the cytoplasmic side and lead to the outward movement of the TM6 helix, a hallmark for GPCR activation. For a given receptor, different ligands can have counteracting effects (e.g. agonizing or antagonizing) or different activation efficacies (e.g. partial, full or supra-activating) ^2^. Moreover, some ligands may preferentially activate the G protein or β-arrestin coupled pathway, known as the biased ligands ^3–5^. Understanding the molecular basis for the diverse ligand efficacies and bias is essential for rational designs of better therapeutics.

Muscarinic acetylcholine receptors (mAChRs) are prototypical GPCRs that respond to the neurotransmitter acetylcholine (ACh) in both the central and peripheral nervous systems ^6^. They are also attractive drug targets for treatment of Alzheimer’s disease, Parkinson’s disease, schizophrenia and cardiovascular diseases ^7–9^. In particular, the M2R subtype has been extensively studied by structural, functional and computational methods, and a handful of small molecule ligands that bind M2R with varying efficacy profiles have been developed ^10–15^. Up to date, active-state structures of M2R stabilized by the supra-agonist iperoxo (Ixo) have been obtained in complex with either the conformational-selective nanobody (Nb9-8) ^11^, the heterotrimeric GoA protein ^12^, or the β-arrestin-1 ^13^. Moreover, cryo-EM structures of M2R-GoA complexes activated by its endogenous agonist ACh have also been solved recently, demonstrating that ACh stabilizes a more heterogeneous M2R-G protein complex than Ixo, exhibiting two distinct G-protein binding conformations ^15^. While it is difficult to directly identify crucial differences in these structures that could explain the distinct efficacies of Ixo and ACh, our previous NMR investigations on M2R dynamics in response to different ligands revealed a highly complex energy landscape in which distinct receptor conformations were stabilized by different ligands ^14^. Many of these conformations are transient in nature, and there is currently no straightforward experimental method to directly visualize these conformations.

Herein, we comprehensively conducted all-atom molecular dynamics (MD) simulations on M2R bound to different ligands. We observed that the antagonists (3-quinuclidinyl benzilate, QNB and tiotropium, Tio), the endogenous agonist ACh and the supra-agonist Ixo can stabilize distinct rotamer configurations for several critical aromatic residues in M2R. We also performed simulations starting from the inactive state structure in which the antagonist is replaced by the supra-agonist Ixo. In one of the trajectories, we observed the process of TM6 outward movement, a hallmark of receptor activation. The results reveal a sequential occurrence of conformational changes in the receptor that propagates across the membrane, and highlight the critical role of the W400^6.48^-F396^6.44^ residue pair in the activation. Moreover, we observe that ACh and Ixo stabilize distinct conformational equilibriums of the Y206^5.58^, which is correlated with NMR observations and offers a molecular basis for understanding the differential activation efficacies of the two agonists. Overall, our results demonstrate the essential contributions of aromatic ring flips to the receptor activation and suggest a weak coupling mechanism between the extracellular and cytoplasmic sides mediated by aromatic residue dynamics.

## RESULTS AND DISCUSSION

### All-atom MD simulations of M2R in different functional states

To explore the molecular mechanism of how distinct ligands modulate M2R activity, we systematically carried out all-atom MD simulations of M2R in different functional states, including the apo state, the inactive states when bound to the antagonist QNB or Tio, and the active states when bound to Ixo or ACh in the presence or absence of Nb9-8. In these simulations, the atomic coordinates of the receptor in the inactive state crystal structure (PDB entry 3UON) ^10^ were used to generate the initial conformation for simulation of the apo and antagonist-bound states, whereas the coordinates in the active state crystal structure (PDB entry 4MQS) ^11^ were used for simulation of the agonist-bound states. For each state, three independent MD traces lasting 3 μs were acquired, mounting to a total simulation time of 9 μs. Because these simulations report on conformational fluctuations of M2R in distinct equilibrium states, but do not uncover structural transitions related to the activation (or de-activation) process, we refer to them as ‘equilibrium-state simulations’ in this manuscript.

In hope of tracing the receptor conformational changes during activation, we performed additional MD simulations by using the inactive state crystal structure (PDB entry 3UON) as the starting conformation and replacing the ligand with the supra-agonist Ixo. Different from the equilibrium-state simulations, these simulations were set up to capture the inactive-to-active transition, and were therefore referred to as the ‘activation-process simulations’ in this manuscript. Three independent MD trajectories lasting 3 μs each were acquired, and in one of the traces we observed an outward movement of TM6 of over 10 Å, indicating an inactive-to-active (or active-like) structural transition, which is the first observation of such a transition in MD studies to our knowledge. A summary of all simulations performed in the current study is listed in **Table 1**.

**Table 1.** Summary of the MD simulation setup in this study.

| Remark | State | Ligand | Nanobody | Simulation time | Starting conformation |
| --- | --- | --- | --- | --- | --- |
| Equilibrium state | apo | N/A | N/A | 3μs×3 | 3UON w/o ligand |
| | QNB | QNB | N/A | 3μs×3 | 3UON |
| | Tio | Tio | N/A | 3μs×3 | 3UON + Tio |
| | Ixo | Ixo | N/A | 3μs×3 | 4MQS |
| | Ixo+Nb9-8 | Ixo | Nb9-8 | 3μs×3 | 4MQS |
| | ACh | ACh | N/A | 3μs×3 | 4MQS + ACh |
| | ACh+Nb9-8 | ACh | Nb9-8 | 3μs×3 | 4MQS + ACh |
| Activation process | Ixo*<sup>1</sup> | Ixo | N/A | 3μs×3 | 3UON + Ixo |

<sup>1</sup> “\*” denotes an activation-process simulation starting from an inactive conformation in which the ligand is replaced by the agonist Ixo.

### Aromatic residues show large conformational fluctuations

We analyzed the root mean square fluctuations (RMSF) of all twenty types of amino acid residues in the equilibrium-state simulation trajectories. We observe that aromatic residues (F, Y and W) not only have a relatively high abundance in the M2R sequence, but also exhibit profound structural flexibility (**Fig. 1a**). These aromatic residues are enriched in the extracellular half of the receptor structure, particularly close to the orthosteric ligand binding pocket (**Fig. 1b**), and play important roles in both ligand binding and receptor activation. Moreover, evidence suggested that ring flipping motions of aromatic residues, in particular the symmetric Phe and Tyr sidechains, requires additional volume within the protein and are closely related with local dynamics, causing the protein ‘breathing’ motions ^16^. Therefore, we systematically examine the conformational dynamics of aromatic residues located in three structural regions: the extracellular ligand binding pocket, the transmembrane (TM) core, and the intracellular region.

**Figure 1.**
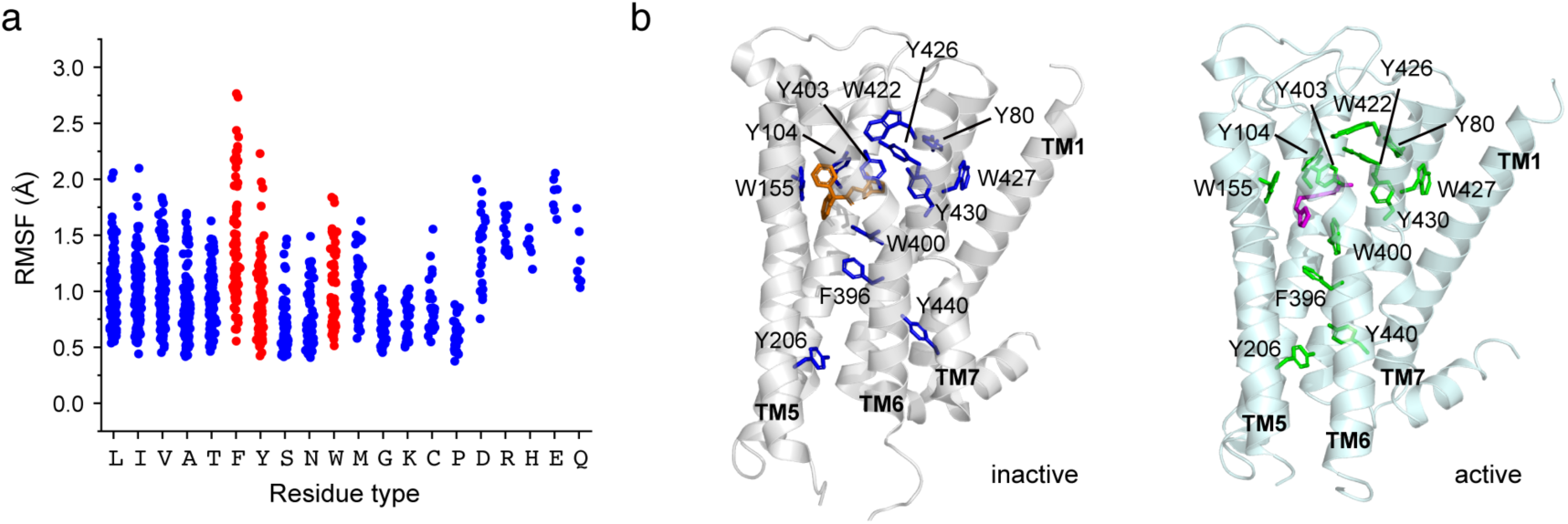
Statistics and location of aromatic residues in M2R. (**a**) RMSF statistics of different amino acids in M2R during the simulations. The residue types are ordered from the highest to lowest according to their abundance in M2R. Each dot corresponds to the RMSF data of an individual residue observed in one equilibrium-state simulation system, averaged among three trajectories. (**b**) Mapping of the locations of important aromatic residues onto the M2R structure in the inactive (PDB entry 3UON) and active (PDB entry 4MQS) states. The ligand QNB and Ixo are colored in orange and magenta, respectively.

To more perceptually describe the conformational fluctuations of the aromatic residues, two analyzing strategies were used. Firstly, we constructed a two-dimensional (2D) statistical heatmap containing information on both sidechain rotation (the χ2 dihedral angle) and translation (distance from the outmost heavy atom in the corresponding aromatic residue to the center of mass of the receptor, denoted as *r* in this manuscript) (see **Methods** and **Supplementary Fig. S1**). Secondly, we calculated the interaction energies between different residue pairs as well as between residues and ligands, so as to provide more quantitative assessments of the protein conformational dynamics.

### Agonist binding stabilizes the aromatic residues at the ligand binding site

A network of aromatic residues surrounding both the orthosteric ligand binding pocket and the allosteric vestibule undergo significant conformational rearrangements between the inactive and active states. These include residues Y104^3.33^, Y403^6.51^ and Y426^7.39^ and Y430^7.43^ that form the “tyrosine cage” surrounding the ligand, as well as the nearby residues Y80^2.61^, W155^4.57^, W422^7.35^ and W427^7.40^. These residues exhibit varying degrees of conformational flexibility in the apo state, whereas binding of antagonists or agonists differentially alters their conformational dynamics (**Fig. 2 & Supplementary Fig. S2-S3**).

**Figure 2.**
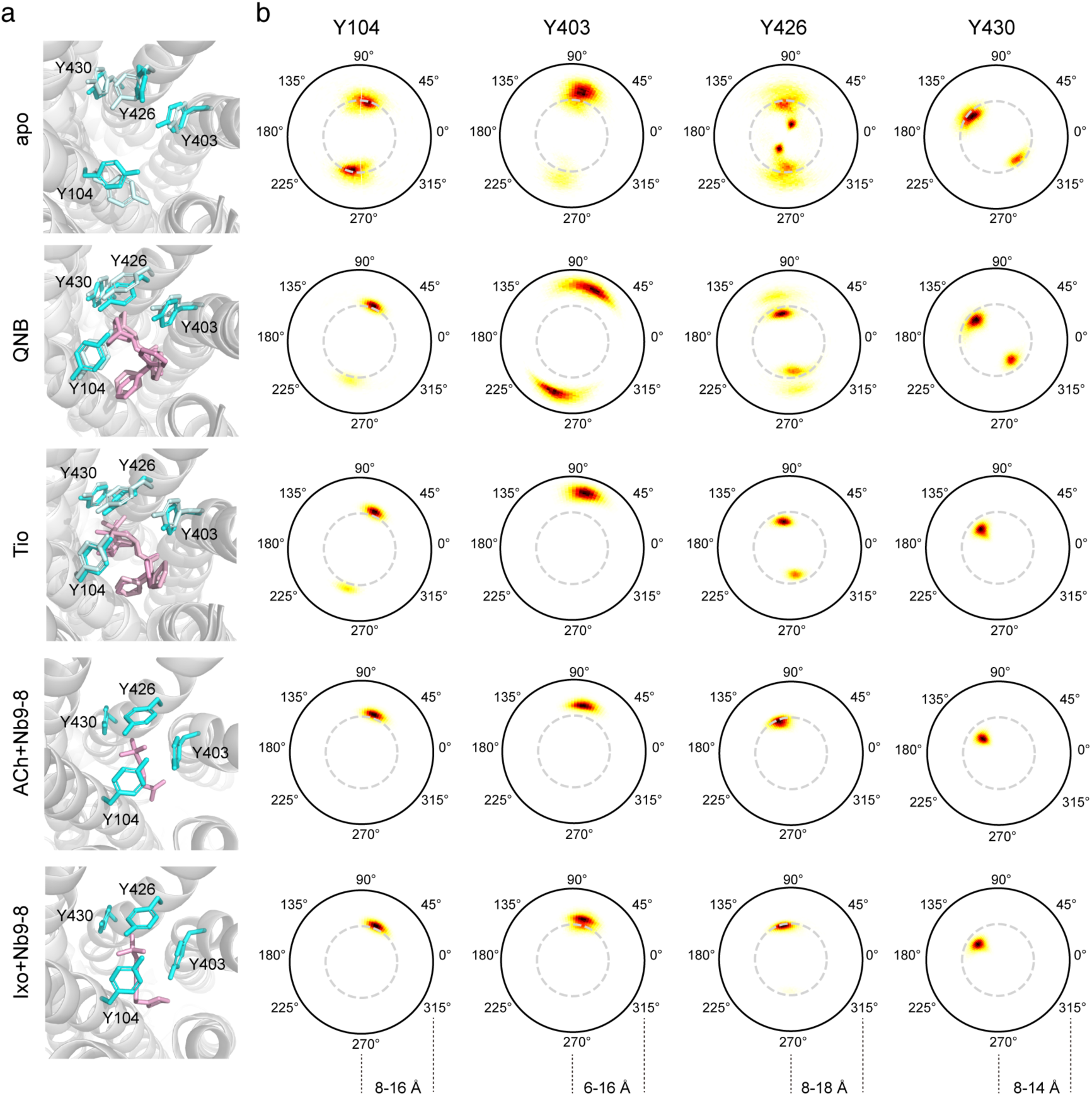
Sidechain conformational equilibrium of the tyrosine cage residues in M2R ligand binding pocket. (**a**) Representative structures showing the sidechain conformations of the tyrosine cage residues in different states. For the apo and antagonist-bound states, two different conformations are shown. Ligands are colored in pink/light pink. (**b**) The 2D heatmaps showing the conformation distributions of the tyrosine cage residues in different states. The angles represent the sidechain χ2. The range of the polar *r* axis is shown at the bottom.

In the apo state, tyrosine residues generally undergo obvious ring flipping events, indicated by the observation of two rotamer conformations with 180° difference in the χ2 angle. Residue Y426^7.39^, in particular, is observed to sample multiple sidechain conformations. In comparison, tryptophan residues undergo less significant ring flipping motions, presumably due to their larger sidechains, but still exhibit obvious conformational heterogeneity. In the simulations of agonist (ACh or Ixo) bound states with or without Nb9-8, all these aromatic residues are stabilized into a unique conformation, and their side chains exhibit no apparent ring flipping motions. Removal of Nb9-8 shows no obvious effects on these residues during the simulation period (**Supplementary Fig. S3**).

Interaction energy analysis suggests that agonist binding significantly enhances the interactions between the Y104^3.33^-Y403^6.51^ and W422^7.35^-Y426^7.39^ residue pairs, while decreasing those between the Y403^6.51^-W422^7.35^ and Y403^6.51^-Y426^7.39^ pairs (**Table S1 & Supplementary Fig. S4**). In particular, the Y104^3.33^-Y403^6.51^ interaction is formed only in the agonist-bound states, but not in the apo or antagonist-bound states.

In contrast to the agonists, antagonist binding does not show much stabilizing effect in the extracellular ligand binding pocket. Most of the tyrosine residues still exhibit obvious ring flipping in the antagonist-bound states, whereas the tryptophan residues (W422^7.35^ and W427^7.40^) adopt two or more different sidechain conformations when bound to QNB or Tio (**Supplementary Fig. S3**). Notably, the larger volumes of the antagonists push the Y403^6.51^ sidechain upward such that it stacks more closely with W422^7.35^ and Y426^7.39^ (**Table S1** & **Supplementary Fig. S4**), and prevent it from forming interaction with Y403^6.51^.

Taken together, agonist binding significantly stabilizes the extracellular ligand binding pocket of M2R, while antagonist binding has limited stabilizing effect and can even induce higher conformational heterogeneity for certain residues. Moreover, the smaller sizes of the agonists facilitate the contact between Y104^3.33^ and Y403^6.51^, thus strengthening the contacts between TM3 and TM6 in the extracellular ends. These observations are consistent with the previous conclusions that the smaller-sized agonists, but not the larger antagonists, can induce a structural contraction in the receptor extracellular region, which in turn allosterically allows for the opening of the intracellular domain.

### Agonist binding enhances aromatic residue dynamics at the TM core

We next examined the conformational fluctuations of two aromatic residues, F396^6.44^ and W400^6.48^, located at the TM core. W400^6.48^, also known as the ‘toggle switch’, is located at the bottom of the orthosteric pocket, while F396^6.44^ is positioned right underneath W400^6.48^. Different from the extracellular region, both residues show enhanced dynamics in the agonist-bound states (**Fig. 3a-d**).

**Figure 3.**
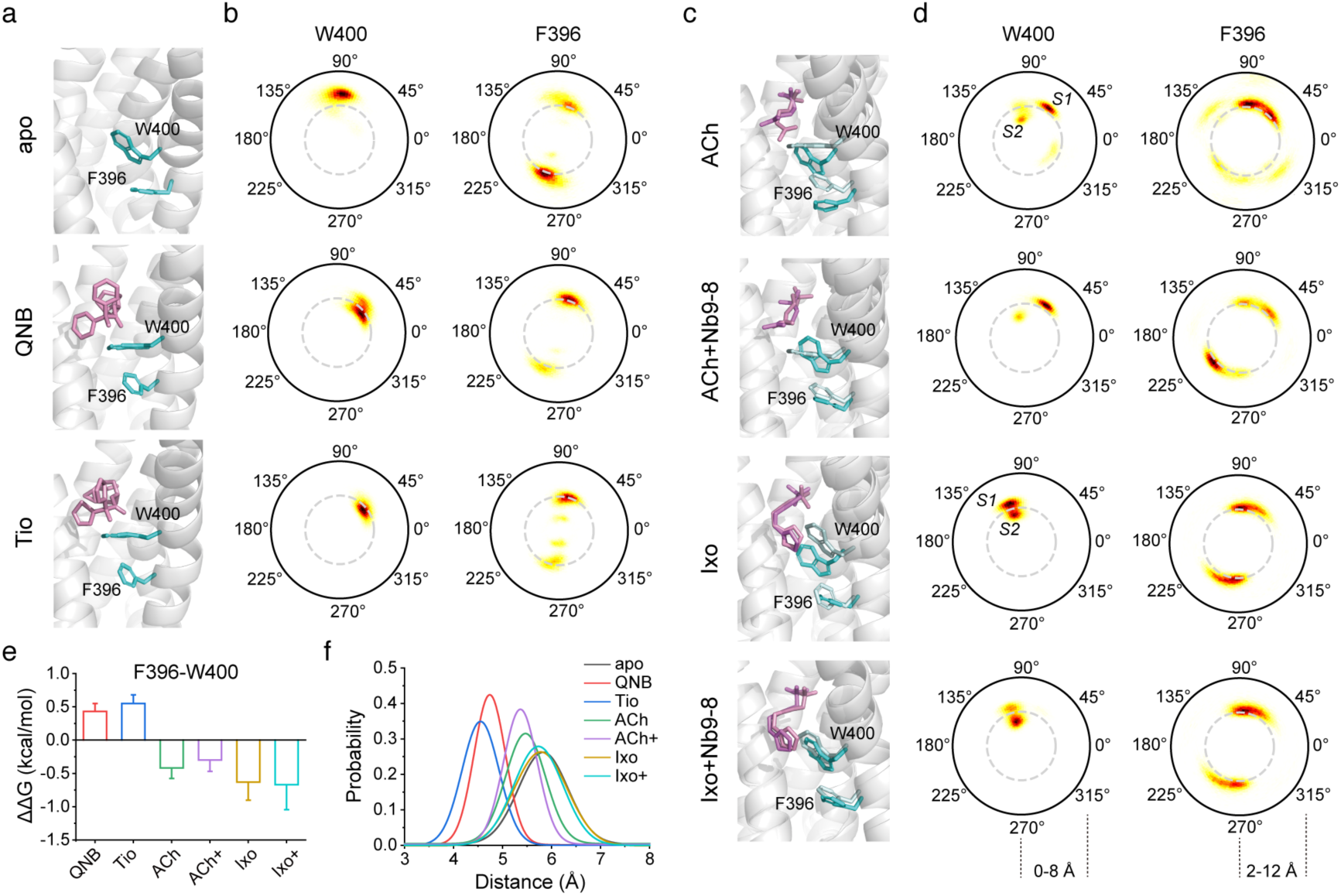
Sidechain conformational equilibrium of aromatic residues at the TM core. (**a,c**) Representative structures showing the sidechain conformations of the aromatic residues in different states. For the agonist-bound states, two different conformations are shown. The W400 and F396 residues are colored in cyan/light cyan. Ligands are colored in pink/light pink. (**b,d**) The 2D heatmaps showing the conformation distributions of W400 and F396 in different states. The angles represent the sidechain χ2. The range of the polar *r* axis is shown at the bottom. (**e**) Changes of interaction energies between F396 and W400. The energy changes are calculated as ΔΔG = ΔGapo-ΔGligand such that a positive ΔΔG value indicates enhanced interaction. **(f)** Distribution of the W400-F396 distance in different states. The distance is calculated between the centers of mass of the aromatic rings of the two residues. In panels e-f, ‘ACh+’ and ‘Ixo+’ denote the ACh+Nb9-8 and Ixo+Nb9-8 states.

The sidechain rotamer configuration of W400^6.48^ is to be strongly affected by ligand binding. In the apo state, it shows a χ2 around 90° with its sidechain oriented upward. Binding of antagonists or agonists can both push W400^6.48^ down towards the receptor core, but stabilize distinct rotamer configurations. On one hand, antagonist (QNB or Tio) binding stabilizes the aromatic plane into a horizontal (relative to the lipid bilayer) orientation with χ2 close to 45°. On the other hand, binding of the native agonist Ach or the supra-agonist Ixo have distinctive effects on W400^6.48^ conformation. In the Ixo-bound state, the aromatic ring of W400^6.48^ is stabilized into two clusters of conformations (S1^Ixo^ and S2^Ixo^), both showing a close-to-vertical orientation with χ2 values around 115° but with different translational movements. In the ACh-bound state, however, the W400^6.48^ sidechain adopts two distinct clusters of conformations, one of which (S1^ACh^) shows χ2 values around 60° and the other (S2^ACh^) is similar to the Ixo-stabilized configuration S2^Ixo^. The different behaviors of these two agonists are likely due to their different end groups. The bulkier isoxazole ring of Ixo can better interact with the aromatic ring of W400^6.48^ and locks it into a close-to-vertical conformation, whereas the smaller and more flexible acetyl group of ACh cannot.

In the case of residue F396^6.44^, significantly elevated conformational dynamics is observed in the agonist-bound states, as indicated by the large variations of the sidechain χ2 angle. The Ixo-bound state, in particular, enables F396^6.44^ to undergo extensive ring flipping motions, as suggested by the equally-populated conformers with 180° difference in the χ2 angle. In contrast, binding of the two antagonists slightly inhibit the ring flipping motions compared to the apo form.

Notably, among the available structures of active-state M2R, the Ixo-activated M2R in complex with either the stabilizing nanobody or with the heterotrimeric G-protein displays a close-to-vertical W400^6.48^ sidechain orientation, which is consistent with the simulation results. In contrast, in the cryo-EM structures of ACh-activated M2R-G-protein complex ^15^, only the close-to-horizontal orientation is observed (**Supplementary Fig. S5**). Because the S2^ACh^ conformation is much lower populated in the MD simulations, information regarding this conformational state is expected to become lost during cryo-EM data averaging. In the M2R-β-arrestin complex structure ^13^, which is activated by Ixo and the positive allosteric modulator LY211960, W400^6.48^ sidechain is also observed to adopt a close-to-vertical orientation. Moreover, the conformational dynamics of F396^6.44^ were also not reflected in these structures, as the residue displays similar sidechain configurations.

### The W400^6.48^-F396^6.44^ interaction is weakened in the agonist-bound states

Because the conformational dynamics of these two aromatic residues are likely to be coupled, we further examined the W400^6.48^-ligand and W400^6.48^-F396^6.44^ interaction energies (**Supplementary Table S2**). The results show that both the two antagonists and Ixo interact stronger with W400^6.48^ compared to the native ligand ACh. However, the interaction between the W400^6.48^-F396^6.44^ residue pair is enhanced upon antagonist binding, and is reduced when bound to the agonists (**Fig. 3e**). Interestingly, the decrease of the W400^6.48^-F396^6.44^ interaction energy is most dramatic upon Ixo binding.

Examination of the structures can provide some clues to these observations. Antagonist binding not only pushes the W400^6.48^ sidechain deeper into the TM core, but also stabilizes a horizontally-oriented W400^6.48^ configuration that allows the large indole plane to form stacking interaction with F396^6.44^. As a consequence, the distance between W400^6.48^ and F396^6.44^ aromatic rings is significantly shortened to below 5 Å (**Fig. 3f**), indicating strong stacking interaction between the two residues. In contrast, although Ixo also show strong interaction with W400^6.48^, it stabilizes the indole ring into a close-to-vertical orientation. This configuration offers limited surface to stabilize the stacking interaction with F396^6.44^, and thus results in decreased W400^6.48^-F396^6.44^ interaction energy and higher F396^6.44^ flexibility. In the case of ACh binding, the S2^ACh^ conformation is similar to S2^Ixo^, while the S1^ACh^ conformation shows a rotamer angle more similar to the antagonist-bound state but is located farther away from the receptor core (**Fig. 3c-d**). Both conformational states lead to weakened interactions with F396^6.44^ (**Supplementary Table S3**), although to a lesser extent compared to the Ixo-bound state. The weakened W400^6.48^-F396^6.44^ interaction may enable F396^6.44^ to exhibit increased sidechain dynamics, including ring flipping motions. Because F396^6.44^ is located at the starting position of TM6 where the outward movement occurs upon activation, we speculate that its enhanced dynamics may have important contribution to receptor activation.

Because F396^6.44^ and W400^6.48^ residues are highly conserved among many Family A GPCRs, we carried out additional MD simulations to verify whether the weakening of W400^6.48^-F396^6.44^ interaction can also be observed in other receptors. The β2AR ^17,18^, A2AR^19,20^ and M1R ^12,21^ receptors, for which both inactive and active structures are available, were chosen. Similar to M2R, three independent simulations lasting 3 μs were conducted for the inactive and active states of these receptors (**Supplementary Table S4**). The results demonstrate that the interaction energies of the corresponding W^6.48^-F^6.44^ residue pair is significantly higher in the inactive state compared to the active state for all the examined receptor systems (**Supplementary Table S5**), suggesting that the weakened W^6.48^-F^6.44^ interaction may be a common feature for the activation of different GPCR members.

### Three sequential steps during Ixo-induced M2R activation

The above observations suggest that weakened W400^6.48^-F396^6.44^ interaction is correlated with M2R activation. However, these results are obtained from the equilibrium-state simulations, and it is unclear whether the weakened interaction is the cause or simply the result of receptor activation since these simulations start from the already-activated conformation. We therefore conducted the ‘activation-process simulations’, which starts from the inactive-state conformation with the ligand replaced by Ixo (**Table 1**), and analyzed the trajectory in which an outward movement of TM6 is observed. The results suggest that the M2R activation process can be divided into three sequential steps.

In the first step, the ligand Ixo quickly positions itself to an energetically-favorable conformation, accompanied by the reorientation of the W400^6.48^ indole ring from the horizontal inactive conformation into an active-like conformation such that it forms a relatively stable stacking with the Ixo isoxazole ring (**Fig. 4a**). This process can occur extremely fast, as the W400^6.48^ sidechain is observed to be already reoriented in the first frame (100 ps) of the pre-equilibration period. Accompanying the rotamer change of W400^6.48^, F396^6.44^ gains increased sidechain dynamics (**Fig. 4b**). These changes are not only observed in the trajectory that captures M2R activation, but also in the other two trajectories in which TM6 remains in an inactive conformation within the simulation time (**Supplementary Fig. S6**).

**Figure 4.**
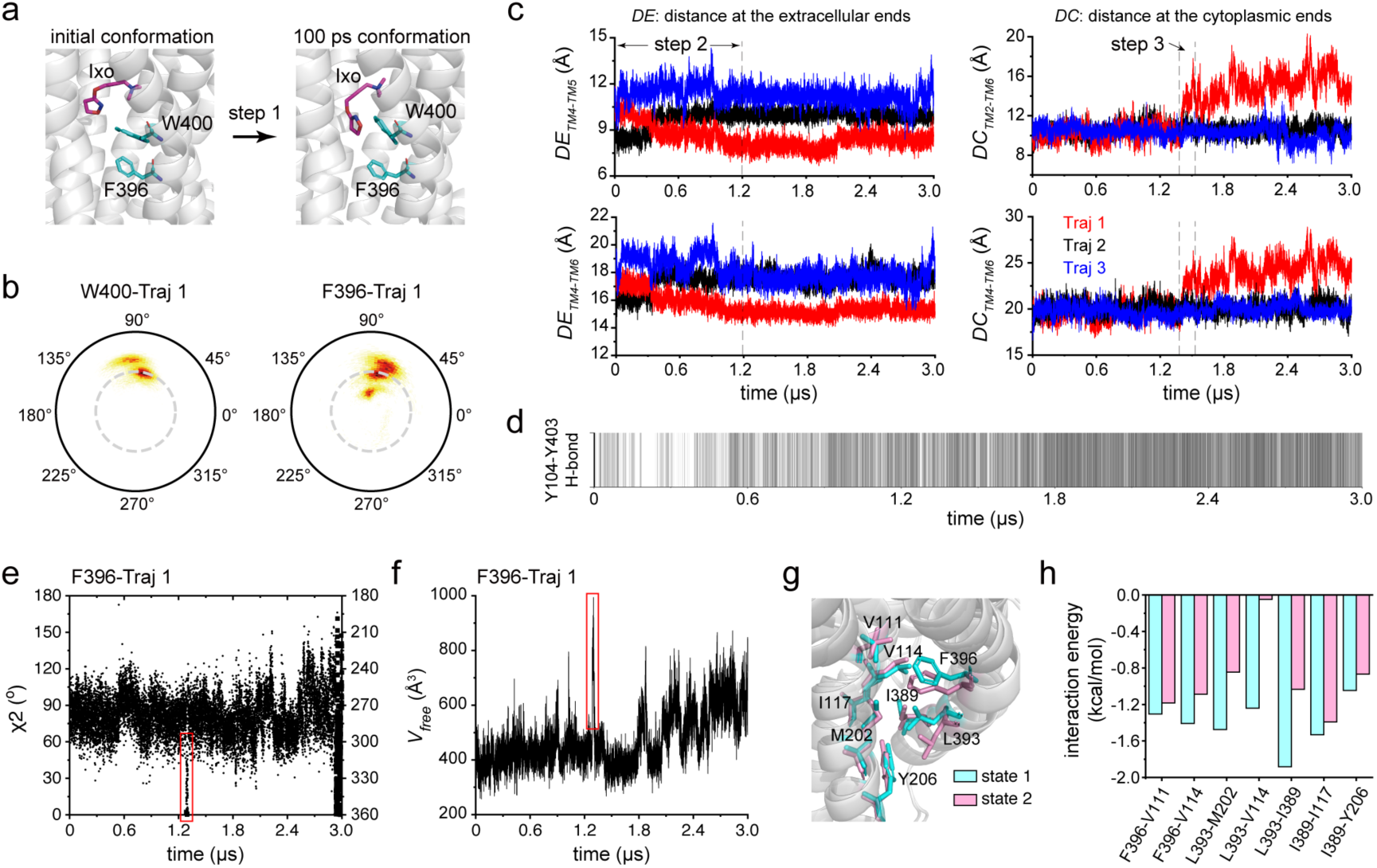
Conformational changes during the activation-process simulation. (**a**) Fast sidechain rotamer change of W400^6.48^ observed in the simulation. (**b**) The 2D heatmaps showing the conformation distributions of W400 and F396 in the activation-process simulation trajectory 1. The angles represent the sidechain χ2. The range of the polar *r* axis is shown at the bottom. (**c**) Changes of inter-helical distances at the extracellular (*DE*) or cytoplasmic (*DC*) ends observed in the three trajectories of activation-process simulations. The distances DE_TM4-TM6_ and DE_TM4-TM5_ were calculated between Cα atoms of W155^4.57^ and V407^6.55^, and those of W155^4.57^ and T190^5.42^. The distances DC_TM2-TM6_ and DC_TM4-TM6_ were calculated between Cα atoms of N58^2.39^ and K384^6.32^, and those of T137^4.39^ and K384^6.32^. (**d**) Time-course analysis of the Y104-Y403 hydrogen bond formation during the activation-process simulation trajectory 1. A single line is drawn for each individual frame if a hydrogen bond is formed. (**e-f**) Time-course changes of the F396 χ2 value (e) and the volume of free space (*V_free_*) (f) during the activation-process simulation trajectory 1. In panel e, the χ2 values in the range of 180-360° are folded back so that the 360° angle returns to the same position as 0°. (**g**) Local structure comparison between two neighboring simulation frames at the time point when the *V_free_* of F396 undergoes sudden increase. (**h**) Comparison of the interaction energies at the TM core between the two frames shown in (g).

In the second step, which corresponds to approximately the first 1.2 μs of the simulation, we observe inward movements of the extracellular ends of several TMs and contraction of the ligand binding pocket in trajectory 1 (**Fig. 4c**). In particular, the extracellular end of TM6 is pulled towards the direction of TM3 and TM4, and the probability of residues Y403^6.51^ and Y104^3.33^ to form a hydrogen bond significantly increases during this period (**Fig. 4d**). Moreover, the extracellular end of TM5 also moves close to TM4, stabilized by interactions between residues N183, V186 and T190 on TM5 and W155, W162 on TM4 (**Supplementary Fig. S7**). In contrast to the extremely fast reorientation of the W400^6.48^ sidechain, the contraction of the extracellular ends of the TMs takes up to 1.2 μs to complete. Note that the extracellular contraction is not observed in trajectories 2 and 3, suggesting that such structural rearrangements are probably prerequisites for the activation of the cytoplasmic side.

In the third step, which occurs at approximately 1.4-1.5 μs, the outward movement of the TM6 cytoplasmic end is observed in trajectory 1 (**Fig. 4c**). Compared to the gradual contraction of the ligand-binding pocket, the outward movement of TM6 occurs significantly faster, taking only about 100 ps to complete. After the time point of 1.5 μs, the receptor maintains in a “TM6-out” conformation with its distance to TM2 (calculated as the distance between residues N58^2.39^ and K384^6.32^) fluctuating between 12 to 20 Å. During this step, we observe that the sidechain dynamics of F396^6.44^ plays an essential role in M2R activation, details of which are analyzed below.

### Analyses of F396^6.44^ dynamics during the activation-process simulation

To understand what might be the molecular trigger of the TM6 outward movement, we closely examined the structural snapshots obtained in the 1.2-1.5 μs period of trajectory 1. In particular, we tracked the time-course of the changes of F396^6.44^ χ2 value and the volume of free space (*Vfree*) surrounding the F396^6.44^ sidechain. Intriguingly, we observe that at approximately 1.3 μs, the rotamer angle of F396^6.44^ starts to deviate from its most comfortable zone (∼ 60-120°) and move into the 0-30° region, which reflects a transient event of the F396^6.44^ sidechain attempting to flip over (**Fig. 4e**). At the same time, the *Vfree* of F396^6.44^, which is normally around 400-500 Å^3^, suddenly increases to close to 1000 Å^3^, forming an obvious spike in the time-dependent profile (**Fig. 4f**). Comparison of two neighboring frames at this time point clearly show that the movement of F396^6.44^ sidechain pushes L393 outward, which alters the hydrophobic interaction network in the TM core (**Fig. 4g**). A number of hydrophobic interactions surrounding the P^5.50^-V^3.40^-F^6.44^ triad become weakened, among which the L393^6.41^-L114^3.43^ interaction is drastically decreased (**Fig. 4h**). Therefore, this observation captures the critical moment when the ring flipping motion of F396^6.44^ disrupts the TM core interactions, offering the opportunity for structural rearrangement and outward movement of TM6. In the subsequent time period (∼ 1.4-1.5 μs), the cytoplasmic half of TM6 is able to break away from the inactive-state position and quickly move out by ∼ 4-5 Å, presumably due to increased kinetic energies in the local region.

Taken together, results from both the equilibrium-state and activation-process simulations support the notion that the W400^6.48^-F396^6.44^ residue pair play essential role in transducing signals across the membrane, and suggest the critical role of F396^6.44^ sidechain dynamics in driving receptor activation.

### ACh and Ixo stabilize distinct conformations in the cytoplasmic cavity

In the cytoplasmic region of the M2R, there are two important aromatic residues Y206^5.58^ and Y440^7.53^, both undergo large scale conformational changes upon activation as revealed by inactive and active-state structures ^10-12^. Although the activation-process simulation captures the TM6 outward movement into an active-like conformation, it is far from enough to allow the visualization of a fully-activated cytoplasmic domain conformation (more discussion on the activation-process simulation is presented in the **Supplementary Text** and **Supplementary Fig. S8**). Therefore, we herein analyze the equilibrium-state simulation results to describe the configurations of these two residues.

While available structures indicate that Y440^7.53^ in the NPxxY motif moves across a distance of ∼7 Å to come close to Y206^5.58^ in the active state, we observe that the Y206^5.58^-Y440^7.53^ contact is unstable in either the ACh- or Ixo-bound state during the equilibrium-state simulations. This observation can be repeated in different simulation setups with or without water molecules in the initial structure to mediate the interaction between the two tyrosines (details are presented in **Supplementary Text**). In brief, while the Y206^5.58^ sidechain is well confined (**Fig. 5a-b**), Y440^7.53^ samples a wide conformational space that deviate from the crystal/cryo-EM structures (**Supplementary Fig. S9**). We speculate that water-mediated Y206^5.58^-Y440^7.53^ contact may be energetically more favorable under freezing or crystalizing conditions, but the NPxxY motif may actually display higher conformational dynamics. Below we focus on analyzing the Y206^5.58^ sidechain configurations.

**Figure 5.**
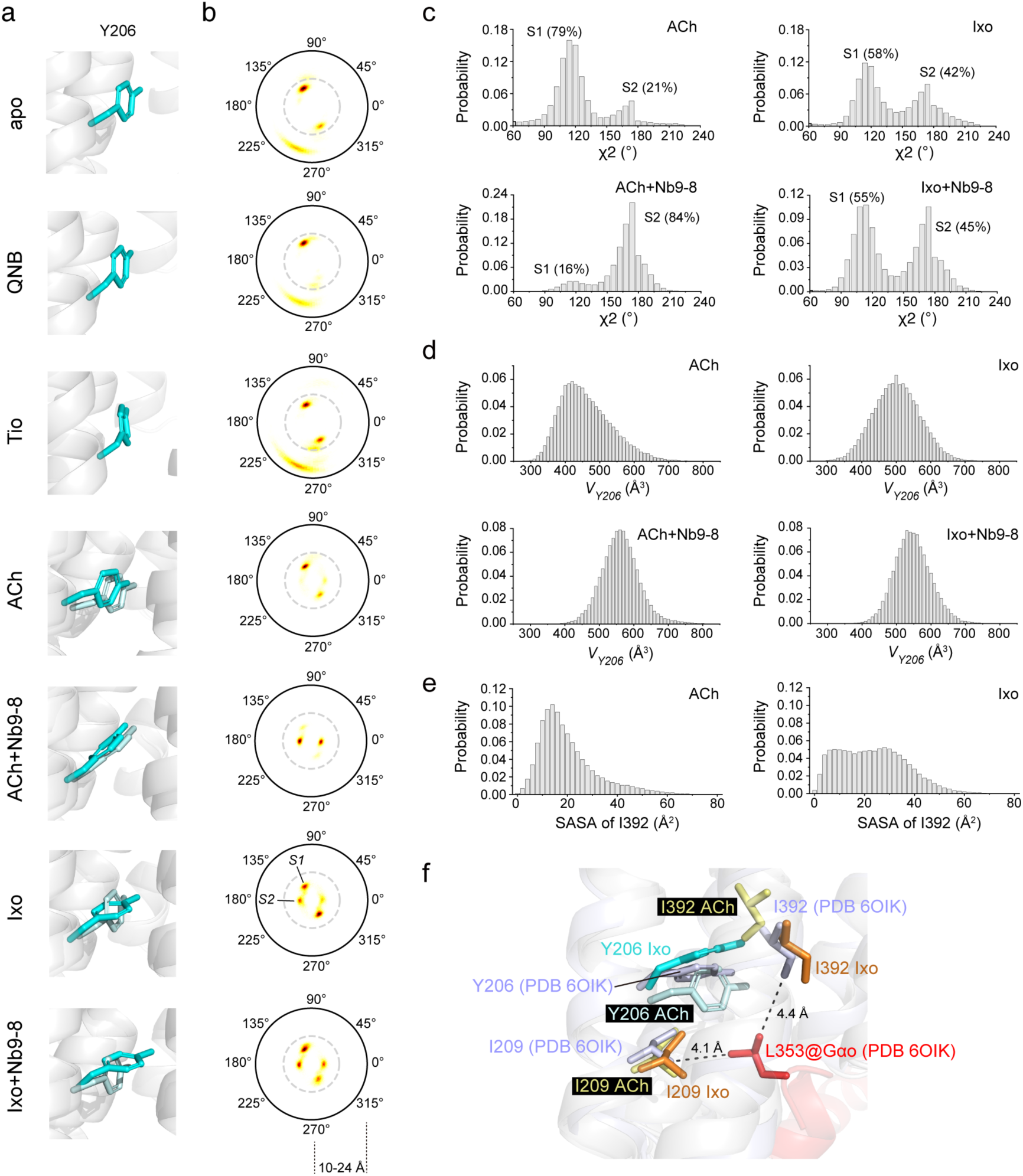
Sidechain conformational equilibrium of residue Y206 at the cytoplasmic side. (**a**) Representative structures showing the sidechain conformations of Y206 in different states. For the agonist-bound states, two different conformations are shown. (**b**) The 2D heatmaps showing the conformation distributions of Y206 in different states. The angles represent the sidechain χ2. The range of the polar *r* axis is shown at the bottom. (**c**) Populations of the S1 and S2 conformations of Y206 in the agonist-bound states. (**d**) Comparison of the free space around residue Y206 (*V_Y206_*) in different agonist-bound states. (**e**) Comparison of the solvent-accessible area (SASA) of I392 in the ACh- and Ixo-bound states. (**f**) Local structure at the cytoplasmic side showing the interactions of residue L353 of Gαo subunit with the I209 and I392 residues of the receptor in the M2R-Ixo-Go complex structure (PDB entry 6OIK). Representative structures of the ACh- and Ixo-bound M2R observed in the simulations are shown to illustrate the different conformations of Y206 and I292 sidechains.

An interesting observation is that, while Ixo binding induces Y206^5.58^ to adopt two rotamer conformations (S1: χ2 ∼120°/300°, S2: χ2 ∼0°/180°), ACh stabilizes Y206^5.58^ into only the S1 conformation. Keeping the nanobody Nb9-8 in the simulation setup slightly modulates the conformational equilibrium of the Ixo-bound state, increasing the S2 population from 40 % to 47 % (**Fig. 5c**). However, the presence of Nb9-8 causes Y206^5.58^ to fully transit into the S2 conformation in the ACh-bound state. Examination of the structural snapshots indicates that the S2 state corresponds to a close-to-horizontal orientation of the Y206^5.58^ ring to the lipid membrane, whereas the aromatic plane is rotated by ∼ 60° towards the vertical orientation in the S1 state (**Fig. 5a**).

Considering the intrinsic shape of the tyrosine ring, we anticipate that the horizontally-oriented S2 configuration of Y206^5.58^ may push the cytoplasmic half of TM6 further away, thus generating a slightly larger intracellular cavity. Indeed, we found that in the absence of a stabilizing nanobody, the volume of the intracellular cavity is smaller in the ACh-bound state (mostly ∼ 400 Å^3^) compared to the Ixo-bound state (∼ 500 Å^3^) (**Fig. 5d**). Binding to Nb9-8 helps stabilize the cavity volume to ∼ 550 Å^3^ for both agonists. Moreover, the cavity size is related to the solvent accessibility of residue I392^6.40^, a key residue that formed intermolecular hydrophobic contacts with L353 (the next to last residue in the C-terminus) of Gαo in the M2R-Go complex ^12,15^ (**Fig. 5e-f**). In the absence of a stabilizing nanobody, the I392^6.40^ sidechain has higher probability of gaining higher solvent exposure in the Ixo-bound state compared to ACh-bound state. Therefore, the Ixo-activated M2R can have a higher probability of initiating interaction with downstream effectors, which provides a structural explanation for the supra efficacy of Ixo.

### Y206^5.58^ sidechain dynamics offer structural basis for previous NMR observations

More intriguingly, the different conformational states of Y206^5.58^ induced by ACh and Ixo binding revealed by the MD simulations correlate well with our previous NMR observations ^14^. The ^13^CH3-labeled methyl group of M202^5.54^, an ideal probe in the cytoplasmic domain, showed distinct NMR spectral changes not only upon binding to ACh and Ixo, but also between binding to ACh alone and binding to both ACh and Nb9-8 (**Supplementary Fig. S10**). Because the anisotropic effect of aromatic ring current has significant impact on local chemical environments, we analyzed the position of the M202^5.54^ methyl carbon atom relative to two nearby aromatic residues, Y206^5.58^ and F396^6.44^, and present the results in the form of 2D heatmap (**Fig. 6a-c**). In this diagram, the radius *r* represents the distance between the ^13^Cε atom of M202^5.54^ methyl group (designated as *C*) and the center of the aromatic ring (designated as *O*), and the angle between the vector 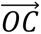 and the aromatic plane is designated as *θ*.

**Figure 6.**
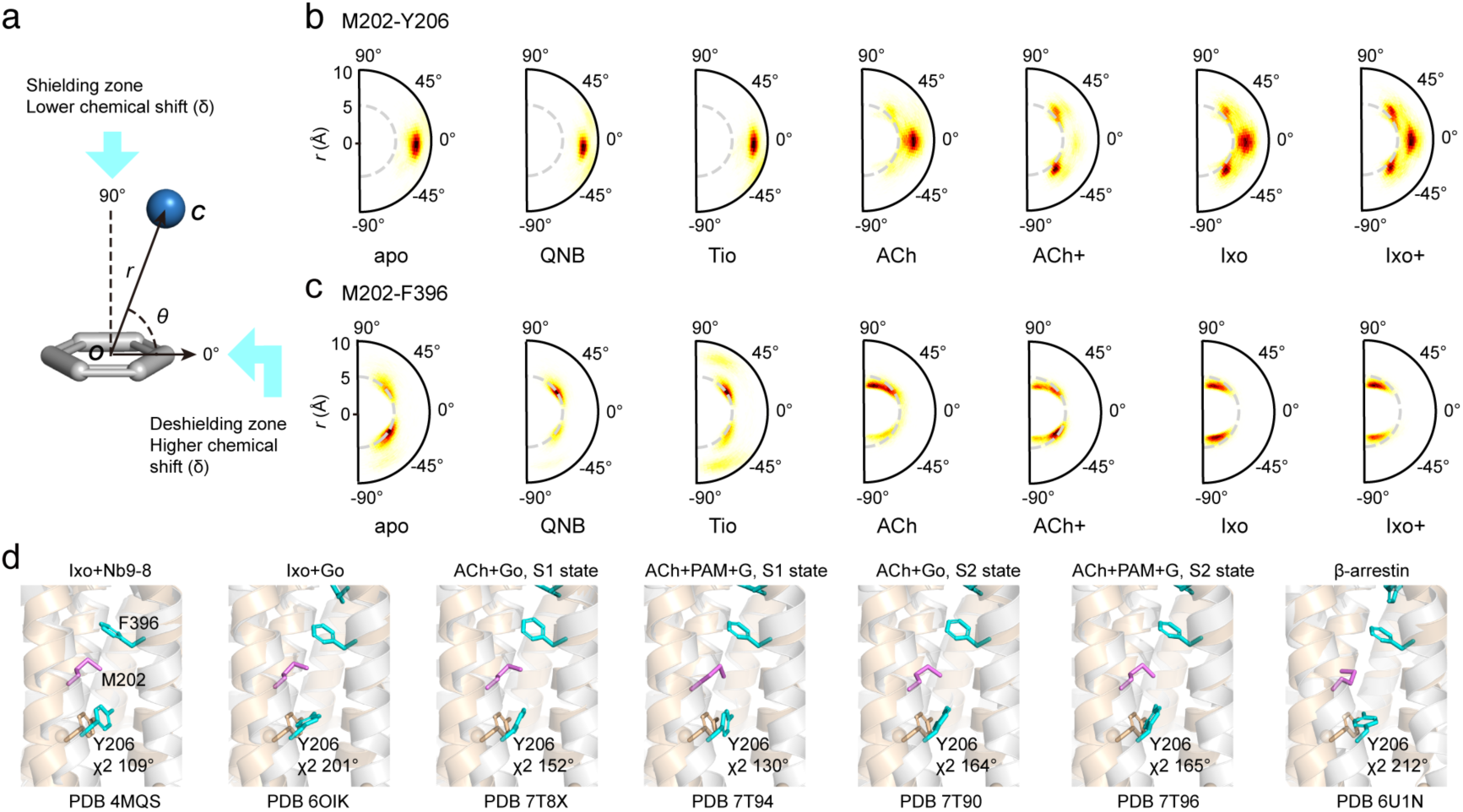
Correlations of MD results with NMR and cryo-EM data. (**a**) Definition of the *θ* and *r* values describing the position of the M202 ^13^Cε atom relative to the aromatic ring. (**b**-**c**) The 2D heatmaps showing the relative position of the M202 ^13^Cε atom to the Y206 (**b**) and F396 (**c**) aromatic rings in different states. (**d**) Different conformations of Y206 sidechain (colored in cyan) observed in the crystal and cryo-EM structures of active M2R. For comparison, the Y206 sidechain in the inactive state structure (PDB entry 3UON) is also shown (colored in wheat).

We found that the *θ* angle between M202^5.54^ and Y206^5.58^ is very close to 0° in the apo and antagonist-bound states, which falls into the deshielding zone with higher chemical shift (δ) values (**Fig. 6a-b**). In the Ixo-bound state, the S1 (close-to-vertical) and S2 (close-to-horizontal) conformations described above correspond to *θ* angles close to 0° and ±45°, which fall into deshielding and shielding zones, respectively. The fast exchanging between S1 and S2 configurations could result in the observation of a single NMR resonance with a smaller ^13^C chemical shift value due to an averaged increased shielding effect. Addition of Nb9-8 to the Ixo-bound state increases the S2 population, causing the ^13^C chemical shift to be further reduced. In contrast, the Y206^5.58^ ring generally adopts the S1 configuration when bound to ACh alone, and therefore the M202^5.54^ ^13^Cε atom remains in the deshielding zone with *θ* angle close to 0°. The small decrease of the *r* value in the ACh-bound state compared to the apo state may account for the slight downfield shift (larger δ) in the ^13^C dimension. When ACh and Nb9-8 are simultaneously bound, the Y206^5.58^ sidechain is stabilized into the S2 conformation, which provides a shielding effect on the M202^5.54^ ^13^Cε atom, causing an opposite change of the ^13^C chemical shift towards the upfield region (smaller δ).

In addition, different from Y206^5.58^, the M202^5.54^ ^13^Cε atom generally falls within the shielding zone of the F396^6.44^ ring in the apo and ligand-bound states (**Fig. 6c**). Agonist-binding stabilizes larger *θ* angles, and Ixo-binding induces *θ* angles closer to ±90° and thus a more significant shielding effect compared to ACh. In the ACh-bound states, the presence of Nb9-8 does not significantly alter the M202^5.54^-F396^6.44^ contact pattern as it does on the M202^5.54^-Y206^5.58^ packing. Taken together, Ixo-induced rotamer changes of both Y206^5.58^ and F396^6.44^ could cause the M202^5.54^ ^13^Cε resonance to move upfield, whereas the distinct rotamer configurations of Y206^5.58^ stabilized by ACh- or ACh/Nb9-8-binding are likely the main contributing factor to the unique NMR spectral changes in the ^13^C dimension.

Moreover, the Hε protons of the M202^5.54^ methyl group show opposite chemical shift changes compared to the carbon atom in the ACh/ACh+Nb9-8 states (**Supplementary Fig. S10**). Because proton is much more sensitive, its chemical shift could be affected by more complex structural factors, e.g. close contacts with nearby oxygen atoms. A detailed discussion of this issue, together with additional analysis on M202^5.54^ χ3 angle-^13^C chemical shift relationship is provided in the **Supplementary Text** and **Supplementary Fig. S11-S12**.

### Y206^5.58^ sidechain dynamics correlates with cryo-EM structures

Available M2R complex structures are also in line with the existence of two different Y206^5.58^ configurations (**Fig. 6d**). When bound to the supra-agonist Ixo, both close-to-vertical (PDB entry 4MQS, M2R-Ixo-Nb9-8 complex) ^11^ and close-to-horizontal (PDB entry 6OIK, M2R-Ixo-Go complex) ^12^ orientations are observed. Among the four available M2R-ACh-Go complex structures (PDB entries 7T8X, 7T90, 7T94 and 7T96) ^15^, conformations closer to the vertical orientation are observed. Note that in the S2 state of the M2R-ACh-Go complex observed in the cryo-EM structure, the Y206^5.58^ rotamer is tilted closer to the horizontal orientation (χ2 values closer to 180°) compared to the S1 state, although the relative orientation between Go and M2R of the S2 state is more similar to the M2R-Ixo-Go complex (note that the S1 and S2 states mentioned here were defined as two states with different M2R-Go packing orientations reported in the previous literature). Additionally, a close-to-horizontal conformation is also observed in the M2R-β-arrestin-1 complex structure (PDB entry 6U1N) ^13^, whereas the Y206^5.58^ sidechain appears more tilted towards the TM3. These static structures, together with our previous NMR data and the current MD simulation results, illustrate the complexity of the intracellular cavity conformation equilibrium regulated by agonist binding.

### Dynamics-driven coupling between the extracellular and the cytoplasmic domains

To further understand why ACh- and Ixo-binding could induce distinct conformations at the cytoplasmic cavity, we carefully examined the structural snapshots in the equilibrium-state simulation trajectories of the ACh- and Ixo-bound M2R without Nb9-8. We observe that in the ACh-bound state, the intracellular region is more likely to deviate from the initial active-state conformation (e.g. the NPxxY motif of TM7 sometimes relaxes back to an inactive-like conformation). Interestingly, we also observe a sliding motion of TM6 along the vertical dimension in some of the simulation frames. As shown in **Fig. 7a**, in the Ixo-bound active state, the horizontally-oriented Y206^5.58^ ring is sandwiched between two hydrophobic residues I389^6.37^ and L393^6.41^, keeping the cytoplasmic end of TM6 in the ‘out’ conformation. Although Y206^5.58^ can flip between the horizontal and vertical orientations, the horizontal conformation is energetically more favorable based on the interaction energies (**Supplementary Table S6**). In the ACh-bound active state, however, while Y206^5.58^ is positioned between the I389^6.37^ and L393^6.41^ residues in most simulation frames, a sub-population of conformers show a vertical sliding of TM6 towards the extracellular region such that Y206^5.58^ no longer stacks in-between the two residues (**Fig. 7b**). This latter conformation, which we call the ‘TM6-up’ conformation for convenience, also shows the characteristics of having the W400^6.48^ rotamer in the horizontal orientation. Thus, a possible scenario is that, because the horizontal W400^6.48^ conformation does not sterically push F396^6.44^ downward as the vertical W400^6.48^ conformation does, the interconversion between the horizontal and vertical conformations of W400^6.48^ induced by ACh binding would make the cytoplasmic part of TM6 prone to move between the ‘TM6-up’ and ‘TM6-down’ conformations. Due to this sliding motion, the Y206^5.58^ sidechain is forced to keep in a vertical orientation.

**Figure 7.**
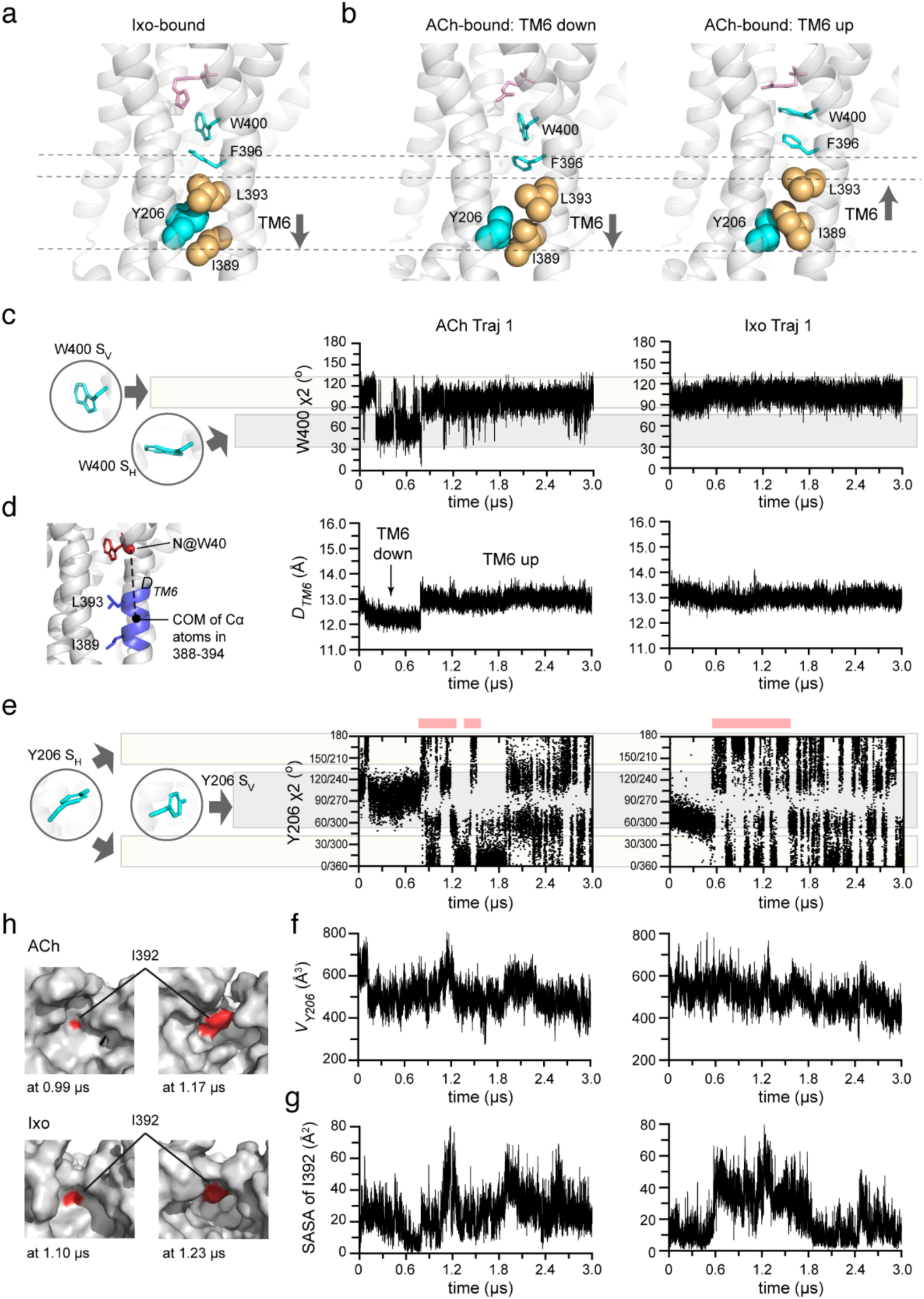
Dynamics-driven coupling between the ligand-binding site and the cytoplasmic cavity. (**a-b**) Sidechain conformation of Y206^5.58^ and its relative position with TM6 in the Ixo-(a) and ACh-bound (b) states observed in the equilibrium state MD simulations. The sidechain atoms of Y206^5.58^, I389^6.37^, L393^6.41^ are shown as spheres, and the sidechains of F396^6.44^ and W400^6.48^ are shown as sticks. The ligands are colored in pink. The ‘up’ and ‘down’ conformations of TM6 observed in the ACh-bound state are shown in (b). (**c**) Time-course changes of W400^6.48^ χ2 angle observed in ACh- and Ixo-bound simulation trajectories. Illustration of the two rotamer conformations corresponding to different χ2 ranges are shown in the left. (**d**) Time-course changes of the TM6 cytoplasmic segment (residues 388-394) in the vertical dimension observed in ACh- and Ixo-bound simulation trajectories. Illustration of the definition of the distance D_TM6_ is shown in the left. (**e**) Time-course changes of Y206^5.58^ χ2 angle observed in ACh- and Ixo-bound simulation trajectories. Pink bars shown on the top indicate the time period where a number of ring flipping events across the horizontal plane are observed. Illustration of the two rotamer conformations corresponding to different χ2 ranges are shown in the left. (**f-g**) Time-course changes of free space around residue Y206^5.58^ (*V_Y206_*) (f) and the solvent-accessible area (SASA) of I392^6.40^ (g) observed in ACh- and Ixo-bound simulation trajectories. (**h**) Representative structural snapshots in the ACh- and Ixo-bound simulation trajectories showing different cavity conformations observed. The structures are shown in surface presentation and the I392^6.40^ residue is colored in red.

Notably, when we examine the time-course changes of the vertical sliding of TM6 together with the rotamer conformations of W400^6.48^ and Y206^5.58^, we found that trajectory #1 of the ACh-bound state well exemplifies this hypothesis (**Fig. 7c-h**). In this trajectory, the W400^6.48^ rotamer changes from the vertical (χ2 ∼ 115°) to horizontal (χ2 ∼ 60°) conformation at the time point of about 0.2 μs, and stayed in the horizontal conformation for a period of ∼ 0.2 μs. During this time, an upward movement of the TM6 cytoplasmic side is observed, and the I389^6.37^, L393^6.41^ and F396^6.44^ residues remained in the ‘up’ conformation until W400^6.48^ flips again into the vertical orientation. These dynamics further affect the rotamer equilibrium of Y206^5.58^, which is stuck in the vertical conformation (χ2 ∼ 100-120°) in the 0.2-0.4 μs time period and becomes free to rotate into the horizontal (χ2 ∼ 0/180°) orientation only when W400^6.48^ adopts a vertical conformation and TM6 moves downwards. During the 0.4-0.7 μs period, we observe the flipping of Y206^5.58^ ring between two chemically identical horizontal configurations (χ2 ∼ 0° and χ2 ∼180°), accompanied by an increase of the *Vfree* around Y206^5.58^ as well as the solvent accessibility of I392^6.40^ to close to 80 Å^2^. Note that in the ACh-bound state, the horizontal orientation of Y206^5.58^ comprises a very small population, and the elevated exposure of I392^6.40^ is also transient. In the 0-0.4 μs time period of trajectory #3 of the ACh-bound state, we also observe continuous flipping events of Y206^5.58^ (with its χ2 angle crossing over the 0°/180° boundaries) associated with enhanced I392^6.40^ exposure (**Supplementary Fig. S13**), suggesting that Y206^5.58^ ring flipping motions may essentially contribute to the modulation of the intracellular cavity conformation.

In the Ixo-bound state, the W400^6.48^ rotamer is stabilized in the vertical conformation and the TM6 is constantly pushed down to allow the Y206^5.58^ sidechain be sandwiched between the I389^6.37^ and L393^6.41^ (**Fig. 7c**). Consequently, the Y206^5.58^ ring has a higher probability of occupying the horizontal conformation and undergoes more frequent ring flipping, which in turn enhances the accessibility of I392^6.40^. The different abilities of ACh and Ixo in stabilizing the W400^6.48^ rotamer into the vertical conformation is transduced into distinct Y206^5.58^ conformational equilibrium and further differentially modulates the I392^6.40^ accessibility (**Fig. 7h**), demonstrating a dynamics-mediated coupling between the orthosteric ligand binding pocket and the intracellular effector-interacting cavity.

### A hypothesized activation mechanism of M2R regulated by aromatic ring motions

Based on the above results, we propose an activation mechanism of M2R in which aromatic ring motions play an essential role (**Fig. 8**). Firstly, agonist-binding induces contraction in the extracellular domain, modulates the conformational equilibrium of the toggle switch residue W400^6.48^ and weakens the W400^6.48^-F396^6.44^ interaction. Among the different sidechain conformations W400^6.48^ might adopt, the close-to-vertical orientation is crucial for receptor activation as is the ring flipping motions of F396^6.44^. The vertically-oriented W400^6.48^ sidechain pushes F396^6.44^ downwards and poses a steric hinderance such that when F396^6.44^ attempts to complete a ring flipping event, it can only gain additional space by pushing away residues located beneath, e.g. L393^6.41^. The motions of F396^6.44^ significantly perturb the TM core interactions and allow the cytoplasmic segment of TM6 to move out. In the ‘TM6-out’ activated state, the Y206^5.58^ conformation equilibrium is modulated by a weak allosteric coupling with the W400^6.48^ configuration. Here again the close-to-vertical orientation of W400^6.48^ is essential to stabilize the stacking of Y206^5.58^ between residues I389^6.37^ and L393^6.41^, and facilitates the Y206^5.58^ ring to undergo ring flipping motions as well as exchange between the horizontal and vertical rotamer configurations. The ring flipping motion of Y206^5.58^ and its probability in occupying the horizontal configuration further modulates the intracellular cavity size and the exposure of the essential I392^6.40^ residue for interaction with downstream effector proteins.

**Figure 8.**
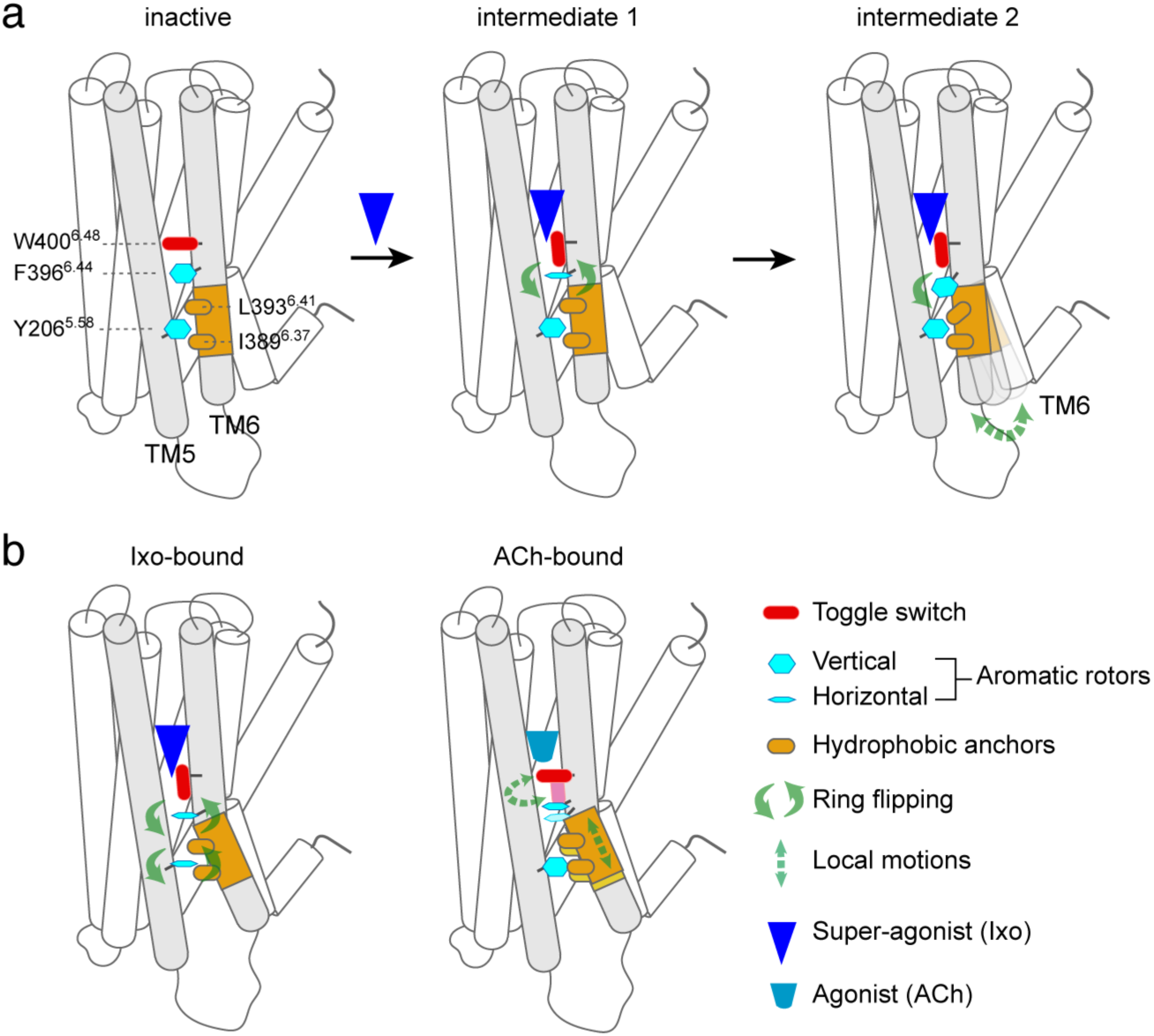
A hypothesized mechanism of M2R activation regulated by aromatic ring motions. (**a**) Schematic illustration showing the ligand-induced activation of M2R regulated by the W400^6.48^ toggle switch and the ring flipping motions of F396^6.44^. (**b**) Schematic illustration showing the weak coupling between the W400^6.48^ conformation equilibrium with the cytoplasmic cavity conformation.

### Conclusions

Our findings offer a structural dynamics basis for understanding the molecular mechanism governing GPCR activation and ligand efficacies. While the aromatic ring flipping motions have been demonstrated important for protein ‘breathing’ using small model proteins ^16^, our results show that they also have fundamental contributions to the functional activation of GPCRs. Our results reveal three important aromatic residues W400^6.48^, F396^6.44^ and Y206^5.58^ that form a pipeline to transduce ligand efficacy information in the form of conformational dynamics across the membrane to the cytoplasmic side. Our model is in agreement with the common notion that a rotamer change of W^6.48^ is important in receptor activation based on the structures of many activated GPCRs. At the same time, our data provide information on receptor conformational dynamics that are difficult to be extracted from cryo-EM or crystal structures. In particular, cryo-EM structures of ACh-bound M2R in complex with Go protein capture only the horizontal configuration of W400^6.48^ and fail to reflect the existence of the low-populated vertical configuration. Moreover, although two different configurations of Y206^5.58^ can be observed in different complex structures, their coupling to the orthosteric pocket and correlation with ligand efficacies were difficult to establish. Complementary to the cryo-EM/crystallographic methods, our previous NMR studies on M2R dynamics demonstrated at residue-level that ACh induces a more heterogeneous receptor conformation compared to Ixo, and revealed peculiar spectral changes in the cytoplasmic cavity region. However, direct translation of these NMR observations into structural models is still extremely challenging. The current MD simulation study comprehensively unravels the distinct conformational dynamics of M2R induced by the binding of different ligands, and offers a structural explanation of the previous experimental results, extending our understanding of receptor activation process and ligand efficacies.

In addition, our work also demonstrates that a combined usage of NMR and MD simulation methods can provide rich information on receptor dynamics. Because aromatic residues are enriched in GPCR sequences and present in many critical micro-switch regions, NMR probes (such as methyl groups) introduced at suitable positions have high potentials of detecting intermediate conformational states that are related to aromatic ring motions. Therefore, we anticipate future investigations to help us gain deeper mechanistic insights and assist drug development based on receptor conformational dynamics.

## Methods

### Simulation system setup

The MD simulations of M2R were based on the QNB bound crystal structure (PDB: 3UON) ^10^ in the inactive state and Ixo bound crystal structure (4MQS) ^11^ in the active state. For the simulation of apo state, the inactive conformation was used without QNB. For the simulations of ACh bound M2R, the Ixo was removed and the ACh was docked into the active conformation of M2R using AutoDock Vina 1.1.2 software ^22^. The G protein-mimetic nanobody Nb9-8 was removed to simulate with the M2R only system. To investigate the activation process of M2R, we used the crystal structure of the inactivated state (3UON) as the initial conformation, removed the antagonist QNB, and docked the Ixo at the orthosteric site. All the simulation systems are listed in Table 1.

For preparing the simulation coordinates, the missing residues in ICL3 region in crystal structure (R216-K234) were modeled using PyMOL (Version 2.3, Schrödinger). The conformation of ICL3 was refined with K234 patched with P377 using Xplor-NIH^23^. To investigate the impact of Nb9-8 on the activation process, we selected conformations of ∼ 1.51 μs during the simulation of M2R activation process. Utilizing the crystal structure containing NB9-8 (4MQS) as a reference, we docked Nb9-8 underneath M2R through superimposition. Further refinements were performed using Xplor-NIH to optimize the positions of ICL3 in M2R and loop region in NB9-8 to prevent atomic clash. This refined structure was used as the initial conformation for subsequent simulations. The membrane environment was prepared for all simulation systems using CHARMM-GUI web server ^24^, and the protein structures were aligned to the Orientation of Protein in Membranes (OPM) ^25^ using PPM 2.0 web server ^26^. Each protein system was inserted into the membrane of dioleoyl-phosphatidylcholine (DOPC) lipids containing ∼70 lipid molecules in upper leaflet and lower leaflet respectively. The water thickness was 20 Å on top and bottom of the system with about 13,000 water molecules for M2R only and 16,000 water molecules for M2R and Nb9-8 complex. The sodium and chlorine ions were added to neutralize the electrostatics at 0.15M NaCl. The final simulation systems were about 80×80×100 Å^3^ for M2R only and 80×80×130 Å^3^ with the addition of Nb9-8. The titratable residues were set in their dominant protonation state at pH 7.0.

### Simulation strategy and analysis

The MD simulations were performed using AMBER22 ^27^ software package with GPU accelerated PMEMD (Particle Mesh Ewald Molecular Dynamics). For all the simulations, the Amber ff19SB force field ^28^ was used to simulate the protein molecules, the lipid21 force filed ^29^ was used for DOPC, GAFF force field ^30^ was used for all the ligands, and TIP3P solvation model was used for water molecules. The simulation systems were first minimized followed by a two-step heating process. First, the systems were heated to 100K in 25 ps with the time step of 1 fs. Then, the heating process increase the temperature to 303K. The protein as well as lipids were restrained with 10.0 kcal mol^-1^ Å^-2^ force constant. After that, the system was equilibrated in the NPT ensemble for 500 ps at 303K with the pressure of 1 bar, with the time step of 2 fs. The production process was also performed in the NPT ensemble for 3μs at 303K with the pressure of 1bar and time step of 2 fs. The time step for production process was also 2 fs to product 15,000 snapshots in each trajectory. For each system, three independent trajectories were performed with random seeds. The SHAKE was applied to constrain the bond length of hydrogen atoms at equilibrium and production stages. The particle mesh Ewald (PME) method ^31^ was applied to treat the long-range electrostatic interactions and non-bonded interactions cutoff was set 10 Å.

The RMSF, dihedral angle, and the corresponding distances were calculated using CPPTRAJ module in AMBER22. The outermost atoms were defined with: OH in Y, CZ in F and CH2 in W. The distances between these outmost atoms and center of mass of the receptor (except for the ICL3 region) were calculated to evaluate the translational movement of aromatic residues. The χ2 dihedral angle were calculated to evaluate rotation of the aromatic sidechain. The 2D heatmap (translational and rotation of aromatic ring) was plot using Matplotlib ^32^ with a bin of 100 snapshots. The interaction energy between the corresponding residues was calculated using the snapshots from last 1μs with MMGBSA ^33^ in AMBER22. The volume around the corresponding residues (F396 and Y206) were calculated using POVME 3.0 ^34^ with the default parameters (distance cut-off of 1.09 Å and grid space of 1.0 Å) for all the snapshots from simulation trajectories. The F396 and Y206 was replaced by glycine respectively to calculate the complete pocket volume. The solvent accessible surface area (SASA) was calculated using NACCESS software ^35^ using PDB files from simulation trajectories. The structure figures were rendered with PyMOL.

## ACKNOWLEDGEMENTS

We thank Prof. Brian. K. Kobilka and Dr. Jun Xu from Stanford University for helpful discussions. This work was supported by the Strategic Priority Research Program of the Chinese Academy of Sciences (XDB0540200) to Z.G. and Y.H., and the Youth Innovation Promotion Association of the Chinese Academy of Sciences (2020329) to Z.G. The authors acknowledge funding from the Joint Laboratory of the National Centers for Magnetic Resonance in Wuhan and in Beijing. Y. H. acknowledges fundings from the Chinese Academy of Sciences and from the Huanghe Talents Plan from Wuhan city.

## AUTHOR CONTRIBUTIONS

C.J. and Y.H. conceived the research. Z.G. performed the MD simulations. Z.G. and Y.H. performed the data analysis and interpretation. X.Z. and M.L. contributed to discussions on data interpretation. Y.H. provided overall supervision. Y.H. wrote the paper with inputs from all authors.

