## Supplementary for "Visualizing the M2 muscarinic acetylcholine receptor activation regulated by aromatic ring dynamics"

This file includes Supplementary Text, Supplementary Figure S1-S13, and Supplementary Table S1-S6.

**Contents:**

**Supplementary Text** s1-s4

Discussion on the effects of Nb9-8 in the activation process of M2R s1

Conformational heterogeneity of Y440$^{7.53}$ observed in MD simulations s2

Discussion on the proton chemical shift of M202$^{5.54}$ methyl group s3

Discussion on M202$^{5.54}$ $\chi 3$ angle distribution and the carbon chemical shift s3

**Supplementary Figure S1-S13** s5-s17

Fig. S1. Illustration of the two parameters used in the 2D heatmap for describing aromatic residue conformations. s5

Fig. S2. Sidechain conformational equilibrium of aromatic residues outside the tyrosine cage in M2R ligand binding pocket. s6

Fig. S3. Sidechain conformational equilibrium of extracellular aromatic residues in the agonist-bound states without Nb9-8. s7

Fig. S4. Interactions at the ligand binding site. s8

Fig. S5. Conformations of W400 and F396 observed in available structures showing two configurations of W400. s9

Fig. S6. The 2D heatmaps showing the conformation distributions of W400 and F396 in the activation-process simulation trajectories 2 and 3. s10

Fig. S7. Conformational changes at the extracellular side in the activation-process simulation trajectory 1. s11

Fig. S8. Effects of Nb9-8 in M2R activation. s12

Fig.S9. 2D heatmaps showing the conformation distributions of Y440 in different states. s13

Fig. S10. NMR spectral changes of the M2R M202 methyl group when bound to ACh or Ixo in the presence or absence of Nb9-8. s14
Fig. S11. Analyses of the M202 methyl proton positions relative to the F396 aromatic ring and the distance between M202 methyl protons with Y206 Oγ atom. s15
Fig. S12. Statistics of the M202 $\chi 3$ angle distribution observed in the MD simulations of different states. s16
Fig. S13. Time-course analysis of M2R conformational dynamics in the #2 and #3 trajectories of the ACh- and Ixo-bound states. s17
**Supplementary Table S1-S6** s18-s20
Table S1. Interaction energies in the extracellular ligand binding pocket in different states. s18
Table S2. Interaction energies between W400-ligand and W400-F396 in different states s19
Table S3. Interaction energies between W400-ligand and W400-F396 in the S1 and S2 conformations when bound to agonist ACh or Ixo. s19
Table S4. Summary of the MD simulation setups for $\beta_2$AR, $A_2$AR and M1R receptors. s19
Table S5. $W^{6.48}$-$F^{6.44}$ interaction energies in different states of $\beta_2$AR, $A_2$AR and M1R. s19
Table S6. Y206-I389 and Y206-L393 interaction energies in different Y206 conformations. s20

### Supplementary Text

#### Discussion on the effects of Nb9-8 in the activation process of M2R

During the activation-process simulation, we capture the TM6 outward movement into an active-like conformation at the time period of ~ 1.4-1.5 μs. However, the cytoplasmic conformation of M2R at this stage is still different from the fully-activated state obtained from cryo-EM or X-ray structures. Moreover, after the time point of about 1.6 μs of the activation-process simulation, we observe that the $Y206^{5.58}$ sidechain flipped out of the TM core and become exposed to the lipids. Because of steric clash with TM6, the $Y206^{5.58}$ sidechain did not flip back again during the rest of the simulation time. We think this may be an off-pathway conformation that cannot lead to functional complex formation with downstream effector proteins or stabilizing nanobodies. We therefore chose a conformation at ~ 1.51 μs time point before the out-flipping event of $Y206^{5.58}$ took place. At this time point, the outward movement of TM6 is ~ 7 Å, and the cytoplasmic cavity is large enough to allow the docking of the Nb9-8 molecule. New simulations (3μs×3) were performed using this structure docked with Nb9-8 as the initial conformation. In two out of the three simulations, $Y206^{5.58}$ maintained an inward-facing orientation while TM6 undergoes further outward movement. In particular, we observe that the TM6 is able to very quickly (in less than 5 ns) open up to reach a TM2-TM6 distance of ~ 20 Å in the presence of Nb9-8 and constantly remain in the 18-19 Å in later stage of the simulation (**Supplementary Fig. S8a-b**). Moreover, we also observe that the cytoplasmic end of TM7 (involving the NPxxY motif) undergoes a conformational change in the presence of Nb9-8 to reach a conformation more similar to the active-state crystal structure (**Supplementary Fig. S8c**). These results demonstrate the important contribution of a downstream interacting partner in inducing the full structural transition of the receptor cytoplasmic domain into the activated state.

### Conformational heterogeneity of Y440$^{7.53}$ observed in MD simulations

During the MD simulations of the active-state M2R bound with either ACh or Ixo, we observe that the the Y206$^{5.58}$-Y440$^{7.53}$ contact is unstable and the Y440$^{7.53}$ sidechain conformation is extremely heterogeneous (**Supplementary Fig. S9**).

Available structures of $\beta_2$AR [1] and μOR [2] suggested that the two conserved tyrosines Y$^{5.58}$ and Y$^{7.53}$ form interactions with each other via a water molecule. However, no such water molecules were observed in the M2R-Ixo-Nb9-8 complex structure [3]. To exclude the possibility that the strong dynamics of Y440$^{7.53}$ observed in the simulations are due to the absence of a stabilizing water molecule, we performed additional simulations starting with initial conformations with water molecules docked into the position to form hydrogen bonds with Y206$^{5.58}$ and Y440$^{7.53}$. The high-resolution μOR crystal structure (PDB entry 5C1M) contains three water molecules close to the two tyrosines, and was used as the reference structure. We built initial structures of ACh- and Ixo-bound M2R (with or without Nb9-8) with all three water molecules added. Simulation results indicate that the water molecules stay near their initial sites for dozens of nanoseconds before moving away. The results are similar between the ACh- and Ixo-bound states, and the presence of Nb9-8 does not have an obvious stabilizing effect.

In addition, we also performed simulations of μOR itself for comparison. The simulations started with μOR structures in which all three water molecules were retained or only one water molecule (the one mediating Y$^{5.58}$-Y$^{7.53}$ interaction) was retained. Simulation results that in the former case, the water molecules stay near their original positions for dozens of nanoseconds, similar to the results of M2R, whereas in the case when only one water molecule is present, it stays near its initial position for only a few nanoseconds. Taken together, these results suggest that while water molecules may have a stabilizing effect on the active conformation in the cytoplasmic side, their interactions with the tyrosine residues are highly dynamic. Therefore, the conformational heterogeneity of Y440$^{7.53}$ observed in the simulation result is not due to the absence of stabilizing water molecules, but more likely reflect the intrinsic dynamics of TM7.

### Discussion on the proton chemical shift of M202$^{5.54}$ methyl group

The Hε protons of the M202$^{5.54}$ methyl group show opposite chemical shift changes compared to the carbon atom in the ACh/ACh+Nb9-8 states. The reason for this may be that protons are more sensitive than carbon and could be affected by more complex factors. For one thing, the averaged positions of the methyl protons are located farther away from the Y206$^{5.58}$ ring (generally larger than 5 Å), and therefore the Y206$^{5.58}$ ring effect may have less effects on the Hε chemical shifts. The M202$^{5.54}$ methyl protons may be more significantly affected by the ring current effects from the F396$^{6.44}$ ring. We observe that in the ACh+ Nb9-8 state, the population shifts towards smaller $\theta$ angles compared to the ACh or the Ixo/Ixo+Nb9-8 states (**Supplementary Fig. S11a**), which may cause a larger shielding effect in the ACh state and contribute to the downfield shift in the proton dimension. For another, the Hε chemical shifts may be more directly affected by its interaction network with nearby residues. For example, we analyzed the distances between the M202$^{5.54}$ Hε atoms and nearby oxygen atoms and observed distinct distributions in the different ligand-binding states. In particular, the M202$^{5.54}$ Hε atoms have a higher tendency to be within 6 Å to the Y206$^{5.58}$ Oγ atom in the ACh+Nb9-8 state compared to all other states (**Supplementary Fig. S11b**), which could also result in a deshielding effect on the Hε atoms in the ACh+Nb9-8 state. These effects, combined with other local conformational differences, may contribute to the downfield shift in the H dimension.

### Discussion on M202$^{5.54}$ $\chi$3 angle distribution and the carbon chemical shift

In a previous NMR study on ACKR3 structural dynamics, the chemical shift changes of M212$^{5.39}$ along the $^{13}$C dimension were attributed to the transition between the *gauche* and *trans* conformations of the methionine sidechain [4]. To evaluate whether the chemical shift of M2R M202$^{5.54}$ methyl group might be affected by a similar mechanism, we statistically analyzed the distribution of the M202$^{5.54}$ $\chi 3$ angle in different states (**Supplementary Fig. S12**). The results suggest that M202$^{5.54}$ adopts a major *gauch* conformation in all states and no significant differences are observed between antagonist- or agonist-binding. Therefore, in the case of M2R M202$^{5.54}$, its sidechain $\chi 3$ rotamer conformation is not sufficient to explain the distinct NMR spectral changes previously observed, and the ring current effect from Y206$^{5.58}$ is more likely the major contributor for the $^{13}$Cε chemical shift changes.

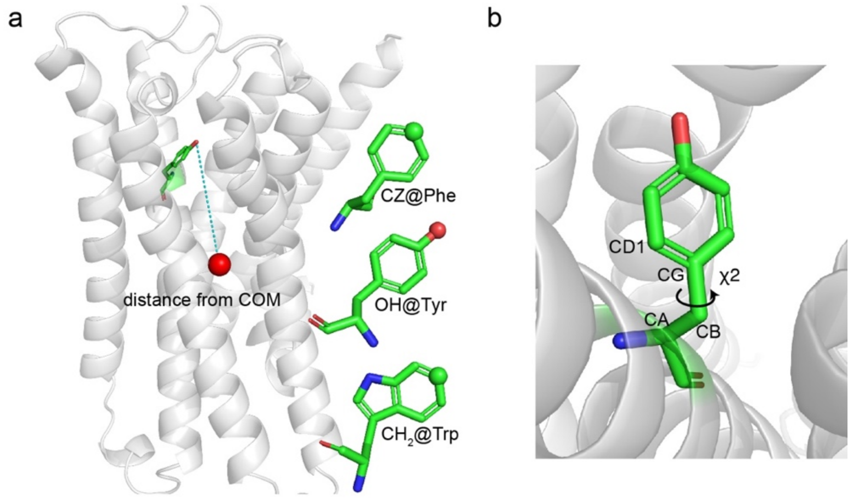


**Figure S1. Illustration of the two parameters used in the 2D heatmap for describing aromatic residue conformations.** (**a**) The parameter *r* is defined as the distance from the outmost heavy atom (shown as a sphere) in the corresponding aromatic residue to the center of mass of the receptor. (**b**) Illustration showing the χ2 dihedral angle of a representative tyrosine residue.

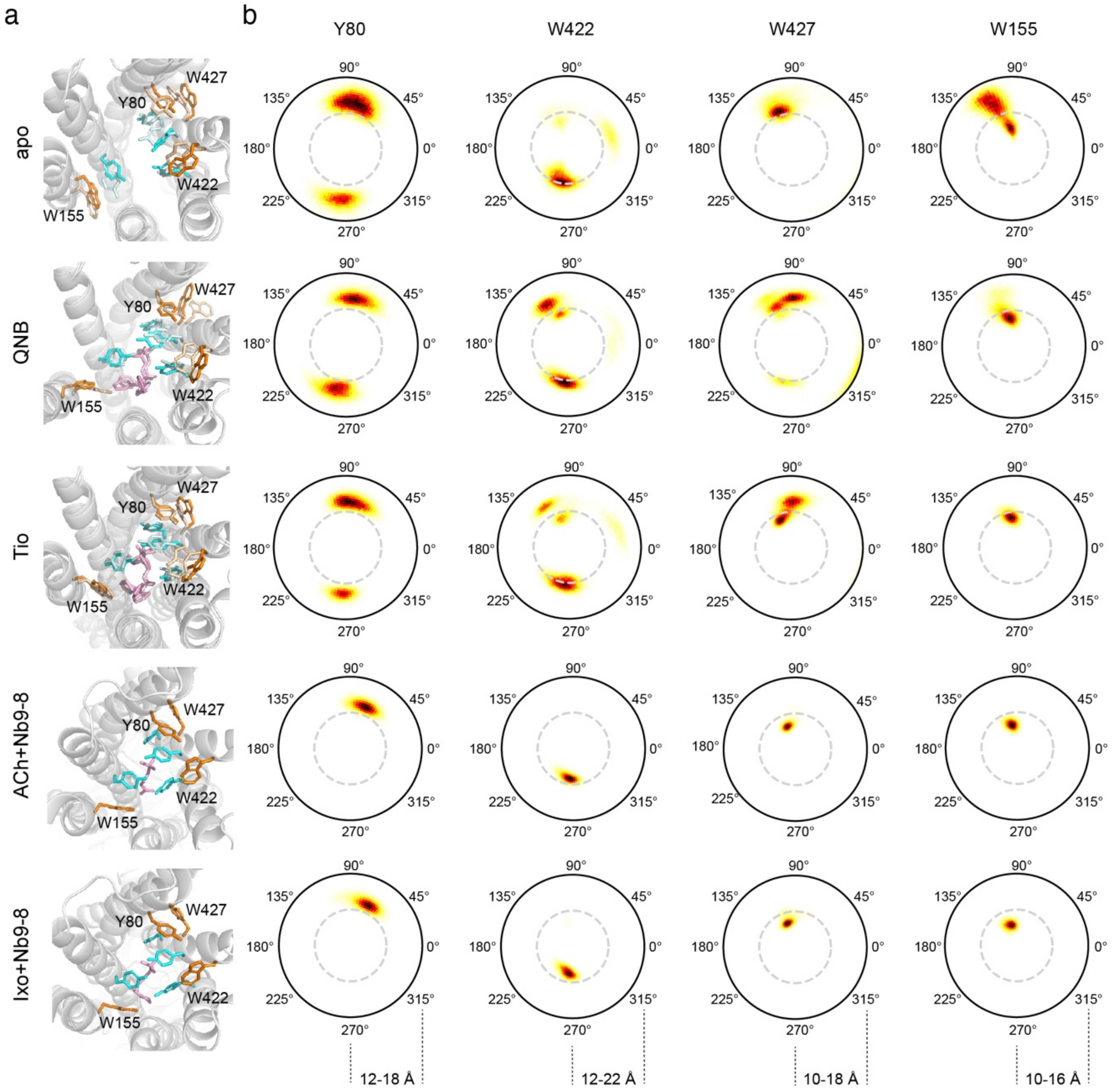


**Figure S2. Sidechain conformational equilibrium of aromatic residues outside the tyrosine cage in M2R ligand binding pocket.** (**a**) Representative structures showing the sidechain conformations of the aromatic residues in different states. For the apo and antagonist-bound states, two different conformations are shown. The Y80, W422, W427 and W155 residues are colored in orange/light orange. The tyrosine cage residues are colored in cyan/light cyan. Ligands are colored in pink/light pink. (**b**) The 2D heatmaps showing the conformation distributions of the aromatic residues in different states. The angles represent the sidechain $\chi 2$. The range of the polar $r$ axis is shown at the bottom.

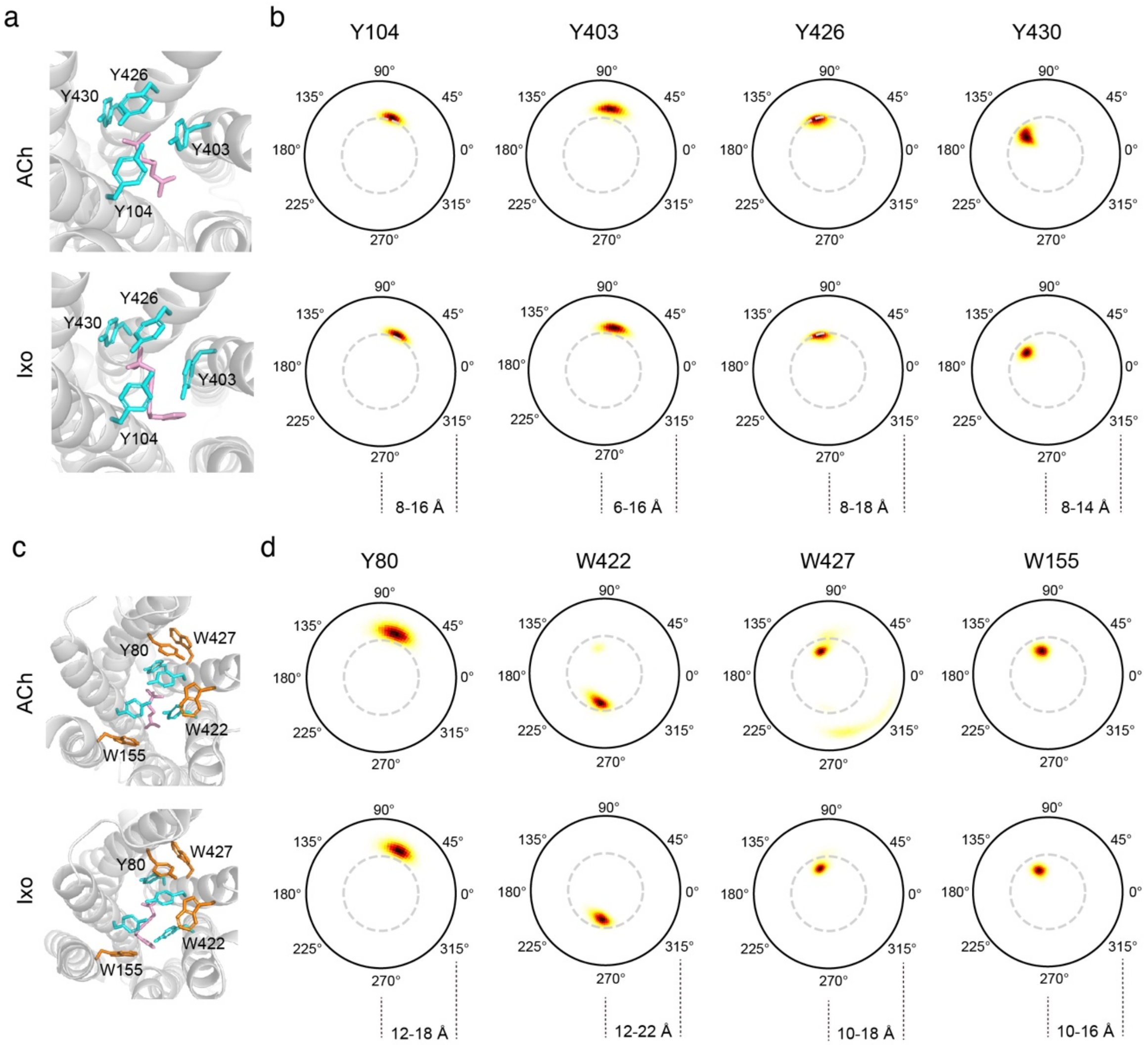


**Figure S3. Sidechain conformational equilibrium of extracellular aromatic residues in the agonist-bound states without Nb9-8.** (**a,c**) Representative structures showing the sidechain conformations of the aromatic residues in different states. (**b,d**) The 2D heatmaps showing the conformation distributions of the aromatic residues in the ACh- or Ixo-bound states in the absence of Nb9-8. The angles represent the sidechain $\chi 2$. The range of the polar $r$ axis is shown at the bottom.

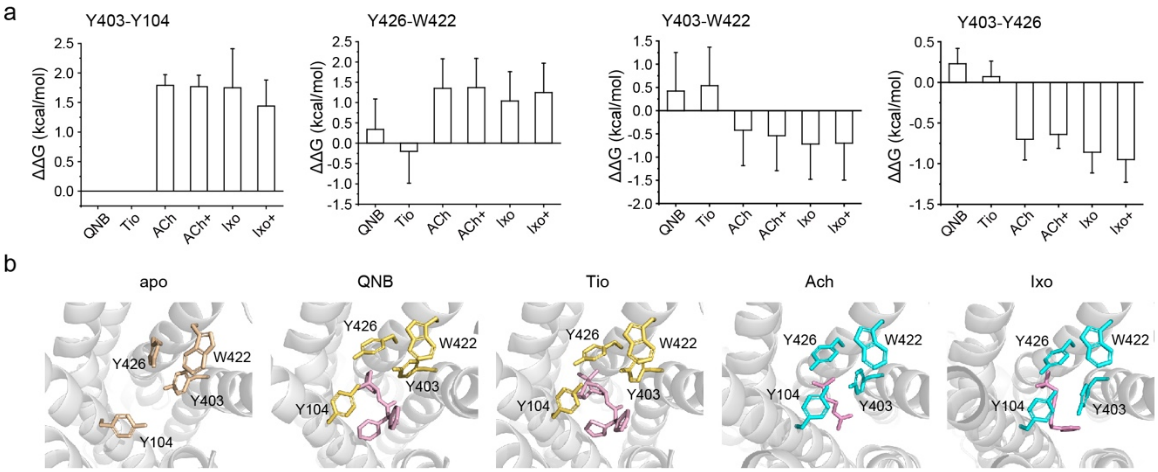


**Figure S4. Interactions at the ligand binding site.** (**a**) Changes of interaction energies between aromatic residue pairs at the extracellular ligand binding site. The energy changes are calculated as $\Delta\Delta G = \Delta G_{apo}-\Delta G_{ligand}$ such that a positive ΔΔG value indicates enhanced binding. In the QNB- and Tio-bound states, the Y104 and Y403 residues do not interact. 'ACh+' and 'Ixo+' denote the ACh+Nb9-8 and Ixo+Nb9-8 states. (**b**) Representative structures showing the aromatic residue contacts at the ligand binding site.

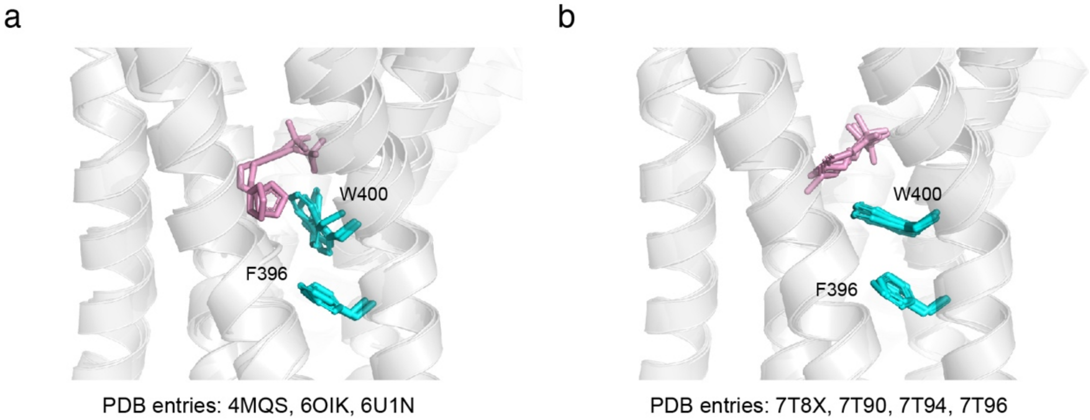


**Figure S5. Conformations of W400 and F396 observed in available structures showing two configurations of W400.** (**a**) Close-to-vertical conformations of W400 observed in the Ixo-bound M2R structures in complex with nanobody Nb9-8 (PDB entry 4MQS), Go protein (PDB entry 6OIK) and β-arrestin (PDB entry 6U1N). (**b**) Close-to-horizontal conformations of W400 observed in the ACh-bound M2R structures in complex with Go protein (PDB entries 7T8X, 7T90, 7T94 and 7T96).

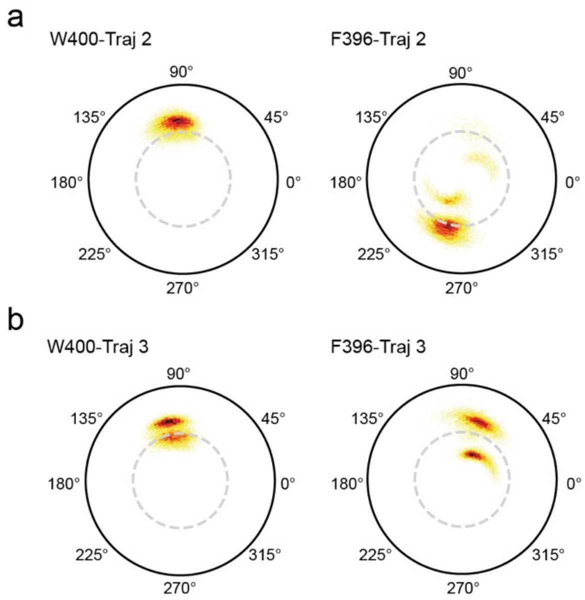


**Figure S6. The 2D heatmaps showing the conformation distributions of W400 and F396 in the activation-process simulation trajectories 2 and 3.**

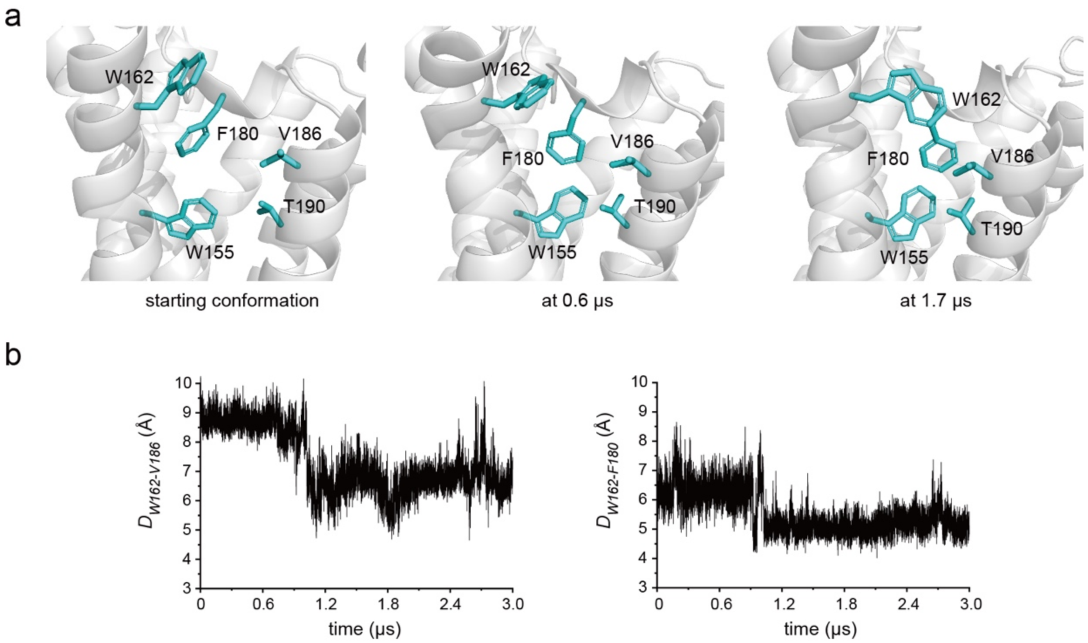


**Figure S7. Conformational changes at the extracellular side in the activation-process simulation trajectory 1.** (**a**) Representative structures showing the interaction changes between TM4 and TM5 at the extracellular side along the simulation trajectory. (**b**) Changes of the W162-V186 and W162-F180 distances during the activation-process simulation trajectory 1. The distances are calculated between the atoms $C\varepsilon_2$ of W162 and $C\beta$ of V186, and between $C\varepsilon_2$ of W162 and $C\gamma$ of F180.

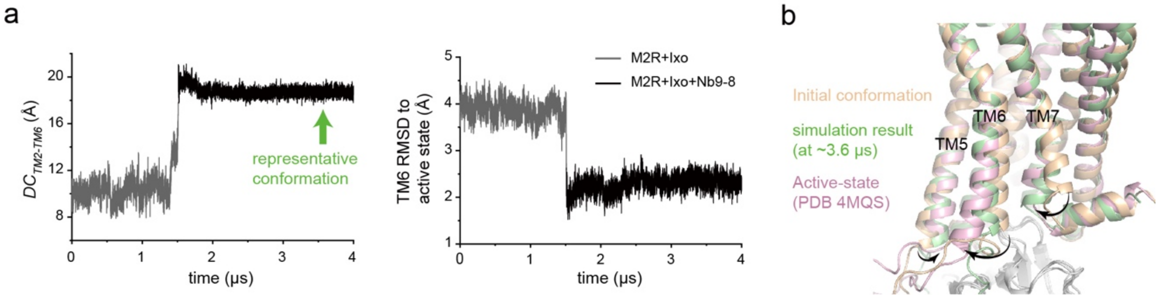


**Figure S8. Effects of Nb9-8 in M2R activation.** (**a**) Conformational changes observed in the activation-process simulation trajectory in which an Nb9-8 molecule was docked into the partially-activated cytoplasmic cavity. The changes of the TM2-TM6 distance at the cytoplasmic side ($DC_{TM2\text{-}TM6}$) is shown in the left panel, and the RMSD of TM6 relative to the active state crystal structure (PDB entry 4MQS) is shown in the right. The grey-colored region corresponds to 0-1.51 μs period of the activation-process simulation trajectory #1 as shown in Figure 5c. The black-colored region corresponds to a simulation trajectory after docking the Nb9-8 molecule into the partially-opened intracellular cavity of M2R. (**b**) Changes of TM6 and TM7 conformations at the cytoplasmic side in the presence of Nb9-8 compared with the active state crystal structure (PDB entry 4MQS).

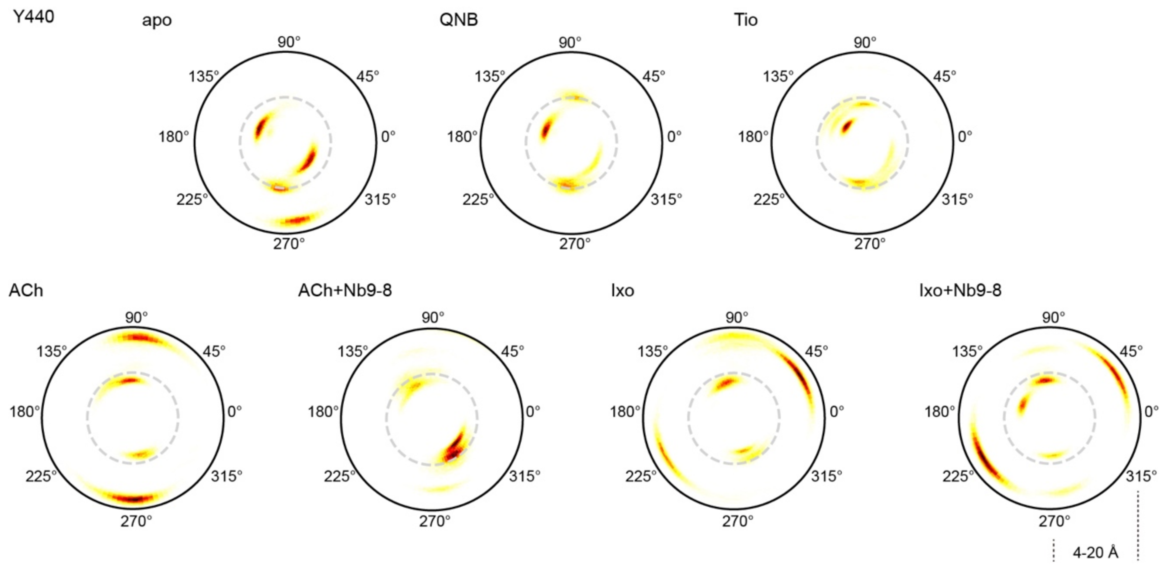


**Figure S9. 2D heatmaps showing the conformation distributions of Y440 in different states.** The angles represent the sidechain χ2. The range of the polar *r* axis is shown.

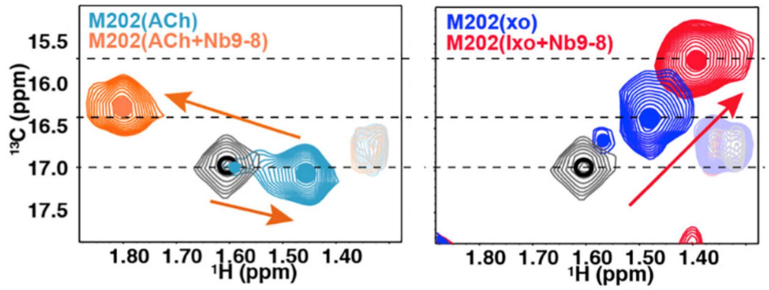


**Figure S10. NMR spectral changes of the M2R M202 methyl group when bound to ACh or Ixo in the presence or absence of Nb9-8.** The figure is reproduced from Figure 6 of previous work (Jun *et al. Mol. Cell.* 2019).

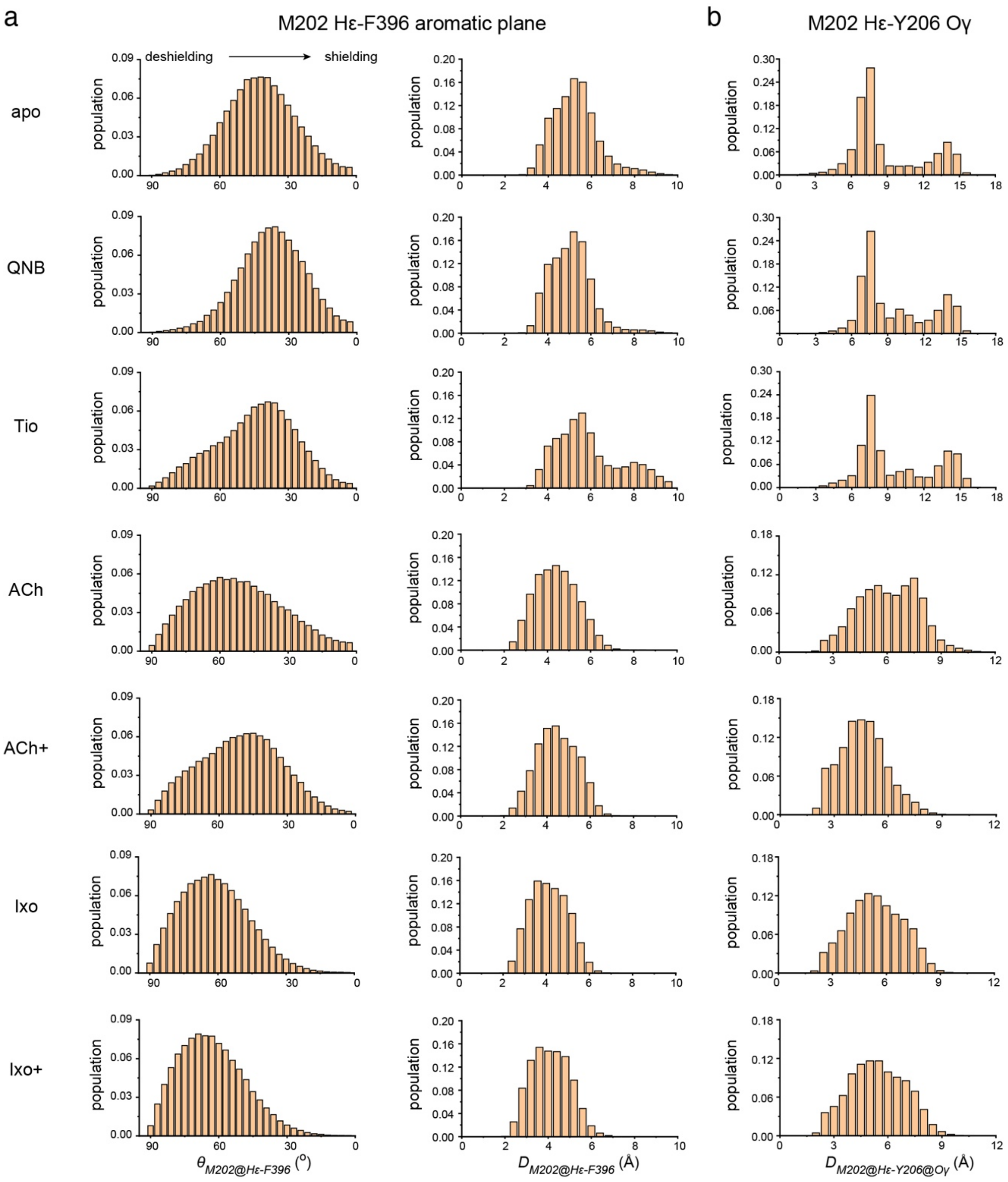


**Figure S11. Analyses of the M202 methyl proton positions relative to the F396 aromatic ring (a) and the distance between M202 methyl protons with Y206 Oγ atom (b).** The $\theta$ angle is defined similarly to that depicted in Figure 6, with the Cε atom replaced by Hε. The distance $D_{M202@H\varepsilon\text{-}F396}$ is defined similarly to the radius *r* in Figure 6, with the Cε atom replaced by Hε.

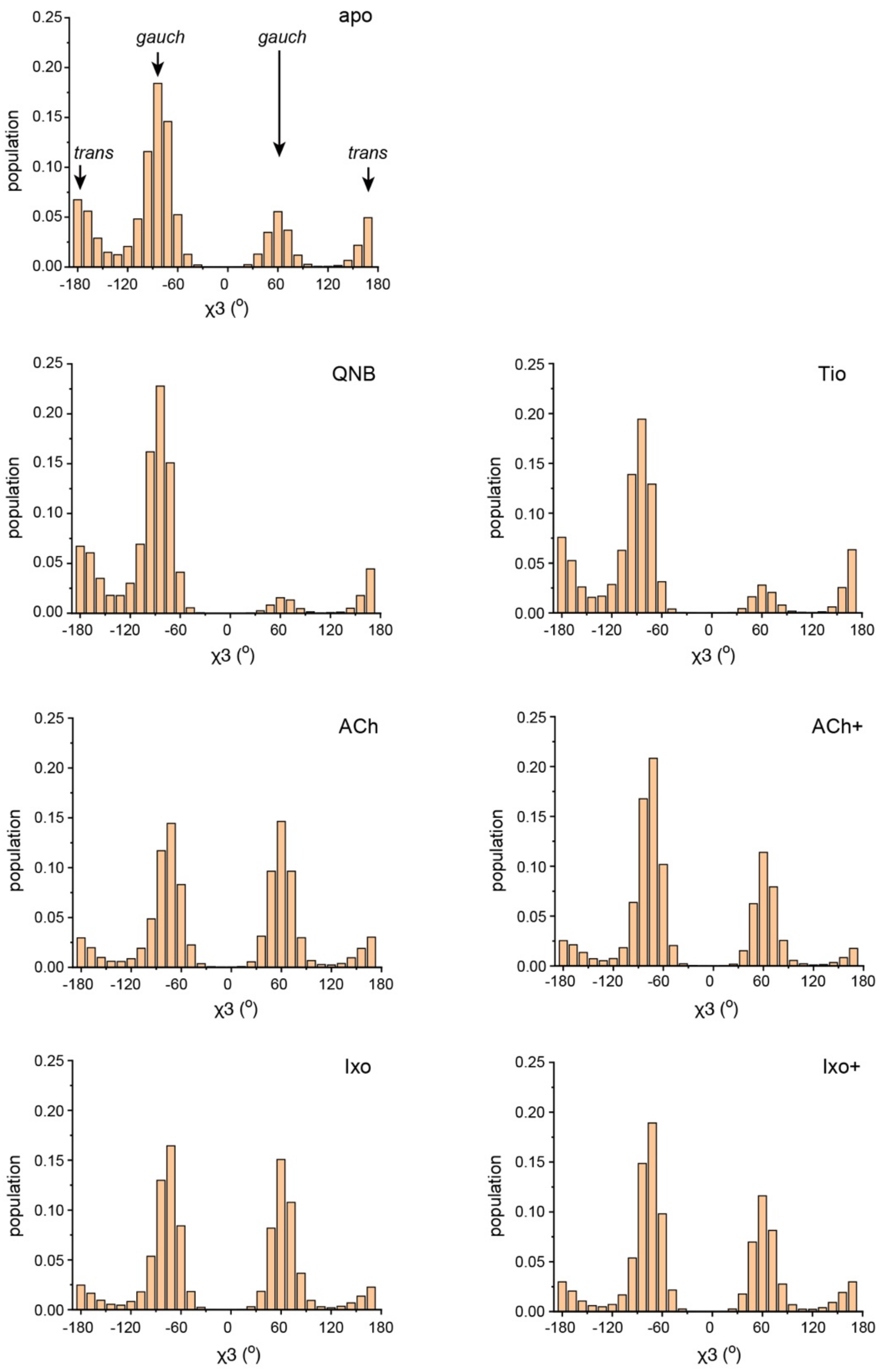


**Figure S12. Statistics of the M202 χ3 angle distribution observed in the MD simulations of different states.** The χ3 angles corresponding to the *gauch* (±60°) and *trans* (±180°) conformations are indicated in the panel of the apo-state.

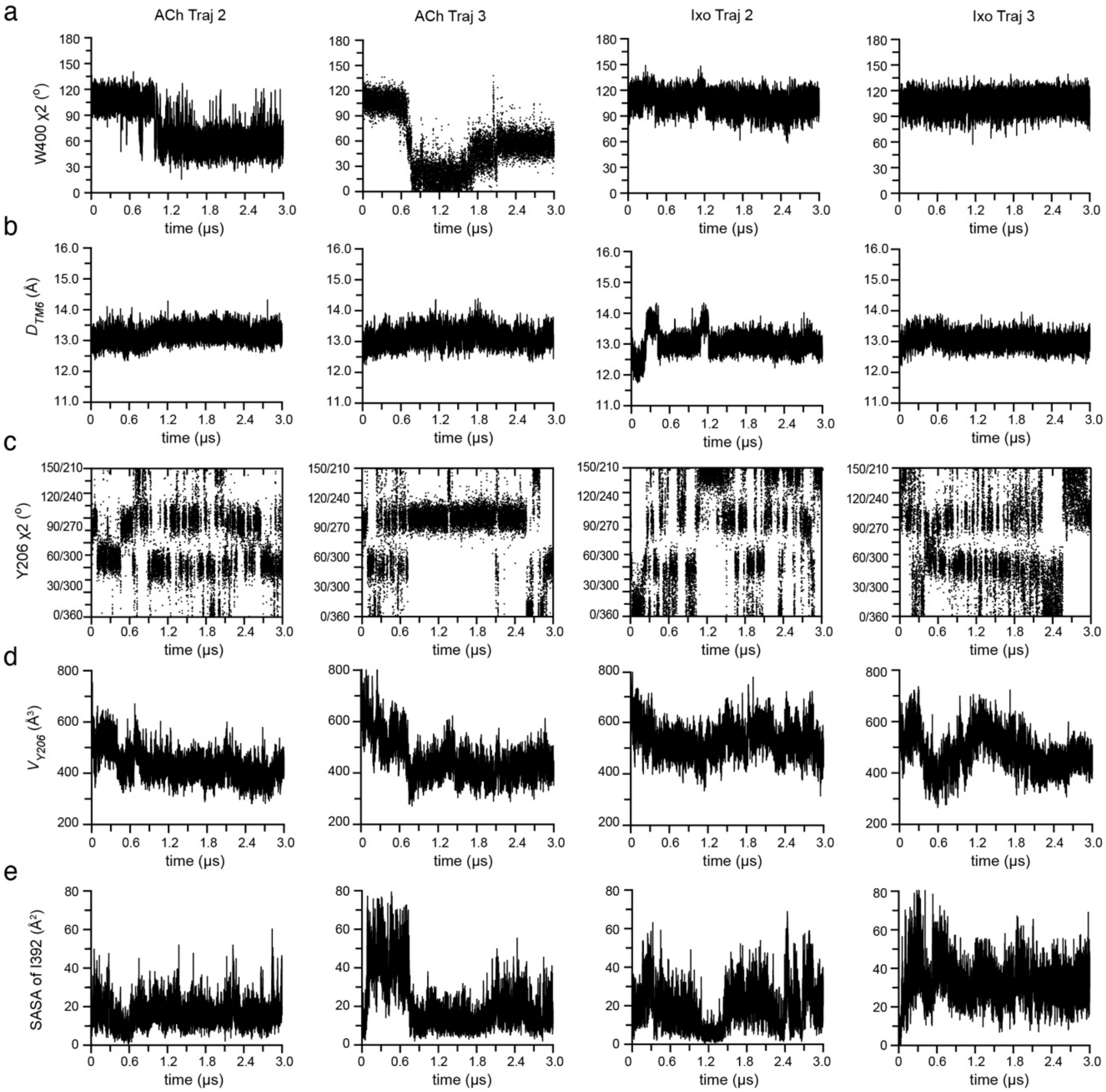


**Figure S13. Time-course analysis of M2R conformational dynamics in the #2 and #3 trajectories of the ACh- and Ixo-bound states.** Time-course changes of W400$^{6.48}$ χ2 angle (**a**), the distance between the TM6 388-394 segment and the backbone nitrogen atom of W400$^{6.48}$ (**b**), the Y206$^{5.58}$ χ2 angle (**c**), the free space around residue Y206$^{5.58}$ ($V_{Y206}$) (**d**) and the solvent-accessible area (SASA) of I392$^{6.40}$ (**e**) observed in ACh- and Ixo-bound simulation trajectories 2 & 3.

**Table S1. Interaction energies in the extracellular ligand binding pocket in different states.**

| | Apo | QNB | Tio | ACh | ACh+ [1] | Ixo | Ixo+ |
|---|---|---|---|---|---|---|---|
| Y104-ligand | N/A | -4.81±0.70 | -5.85±0.30 | -6.16±0.04 | -6.14±0.19 | -5.77±0.21 | -5.73±0.13 |
| Y430-ligand | N/A | -1.85±0.11 | -2.30±0.02 | -2.31±0.28 | -2.17±0.07 | -2.58±0.03 | -2.62±0.12 |
| Y403-ligand | N/A | -3.50±0.54 | -3.53±0.17 | -3.65±0.25 | -3.89±0.22 | -3.83±0.23 | -4.23±0.60 |
| Y426-ligand | N/A | -3.23±0.17 | -5.93±0.26 | -4.76±0.09 | -4.77±0.24 | -5.08±0.18 | -5.01±0.16 |
| Y403-Y104 | N/A | N/A | N/A | -1.79±0.18 | -1.77±0.19 | -1.75±0.06 | -1.44±0.44 |
| Y403-W422 | -2.17±0.72 | -2.59±0.36 | -2.71±0.35 | -1.75±0.14 | -1.63±0.07 | -1.45±0.11 | -1.47±0.27 |
| Y403-Y426 | -2.72±0.17 | -2.95±0.08 | -2.79±0.09 | -2.02±0.19 | -2.08±0.02 | -1.86±0.19 | -1.77±0.22 |
| Y426-W422 | -2.57±0.71 | -2.91±0.24 | -2.37±0.33 | -3.61±0.13 | -3.82±0.13 | -3.92±0.16 | -3.94±0.11 |
| Y430-Y80 | -1.79±0.11 | -1.48±0.05 | -1.28±0.16 | -1.37±0.38 | -1.15±0.01 | -1.25±0.03 | -1.26±0.01 |
| Y430-Y426 | -2.75±0.32 | -1.76±0.11 | -1.42±0.12 | -1.33±0.13 | -1.31±0.04 | -1.32±0.09 | -1.51±0.26 |
| Y430-W427 | -4.04±0.12 | -3.71±0.55 | -4.21±0.63 | -4.26±0.61 | -4.78±0.05 | -4.77±0.12 | -4.68±0.08 |

[1] 'ACh+' and 'Ixo+' denote the ACh+Nb9-8 and Ixo+Nb9-8 states.

**Table S2. Interaction energies between W400-ligand and W400-F396 in different states.**

| | Apo | QNB | Tio | ACh | ACh+ [1] | Ixo | Ixo+ |
|---|---|---|---|---|---|---|---|
| W400-ligand | N/A | -3.64±0.32 | -3.93±0.17 | -2.54±0.62 | -2.61±0.19 | -4.28±0.30 | -4.24±0.39 |
| W400-F396 | -4.48±0.11 | -4.91±0.05 | -5.03±0.07 | -4.06±0.11 | -4.18±0.13 | -3.85±0.25 | -3.81±0.36 |

[1] 'ACh+' and 'Ixo+' denote the ACh+Nb9-8 and Ixo+Nb9-8 states.

**Table S3. Interaction energies between W400-ligand and W400-F396 in the S1 and S2 conformations when bound to agonist ACh or Ixo.**

| | $S1^{ACh}$ | $S2^{ACh}$ | $S1^{Ixo}$ | $S2^{Ixo}$ |
|---|---|---|---|---|
| W400-ligand | -2.57±0.32 | -1.86±0.46 | -4.24±0.41 | -4.47±0.43 |
| W400-F396 | -3.97±0.13 | -4.01±0.21 | -3.95±0.26 | -3.73±0.35 |

**Table S4. Summary of the MD simulation setups for $\beta_2$AR, $A_2$AR and M1R receptors.**

| Receptor | State | Ligand | Simulation time | Starting conformation |
|---|---|---|---|---|
| $\beta_2$AR | inactive | carazolol | 3μs×3 | 2RH1 |
| | active | BI-167107 | 3μs×3 | 3P0G |
| $A_2$AR | inactive | ZM241385 | 3μs×3 | 5IU4 |
| | active | N-Ethyl-5'-Carboxamido Adenosine | 3μs×3 | 5G53 |
| M1R | inactive | Tio | 3μs×3 | 5CXV |
| | active | ACh | 3μs×3 | 6OIJ |

**Table S5. $W^{6.48}$-$F^{6.44}$ interaction energies in different states of $\beta_2$AR, $A_2$AR and M1R.**

| | $\beta_2$AR | $A_2$AR | M1R |
|---|---|---|---|
| Inactive | -4.19±0.11 | -4.20±0.11 | -4.53±0.13 |
| Active | -3.49±0.75 | -3.60±0.20 | -3.76±0.31 |

**Table S6. Y206-I389 and Y206-L393 interaction energies in different Y206 conformations.**

| Y206 rotamer configuration | Residue pair | Interaction energy (kcal/mol) |
|---|---|---|
| Close-to-vertical[1] | Y206-I389 | -2.13±0.32 |
| | Y206-L393 | -1.49±0.32 |
| Close-to-horizontal[2] | Y206-I389 | -3.78±0.42 |
| | Y206-L393 | -1.64±0.36 |

[1] A total of 200 structural snapshots (simulation frames 800-1000) in trajectory 1 of the Ixo-bound state with Y206 $\chi 2$ angles distributed in the range of ~ 60-120° were selected to estimate the averaged interaction energies corresponding to the close-to-vertical sidechain configuration of Y206.

[2] A total of 200 structural snapshots (simulation frames 3800-4000) in trajectory 1 of the Ixo-bound state with Y206 $\chi 2$ angles distributed in the range of ~ 100-180° were selected to estimate the averaged interaction energies corresponding to the close-to-horizontal sidechain configuration of Y206.
